# Ion Mobility-Guided Tandem Mass Spectrometry Imaging Resolves Bis(monoacylglycero)phosphate and Phosphatidylglycerol Isomers in Tissue

**DOI:** 10.64898/2026.08.20.745967

**Authors:** Emanuela Salviati, Fabrizio Merciai, Sandro Montefusco, Aniello Emanuele Giacco, Diego Luis Medina, Pietro Campiglia, Eduardo Sommella

## Abstract

Molecular specificity remains a major challenge in mass spectrometry imaging (MSI), particularly when low-abundance species coexist with structurally related isomers that cannot be distinguished by accurate mass and exhibit similar fragmentation behavior. Bis(monoacylglycero)phosphates (BMPs), lysosomal lipids increasingly implicated in lipid homeostasis and disease, represent a particularly demanding example because they are structural isomers of phosphatidylglycerols (PGs) and display highly similar negative-ion fragmentation. Here, we developed an ion mobility-guided targeted MALDI-MS/MS imaging workflow for direct on-tissue discrimination of endogenous BMP/PG isomeric pairs. Orthogonal HILIC-DDA-PASEF analysis provided accurate-mass, retention-time, fragmentation, and ion-mobility information used to define mobility-constrained precursor coordinates for scheduled MALDI-iPRM-PASEF acquisition. Ion-mobility measurements showed high agreement across ESI-TIMS, MALDI-TIMS, and tissue-based MALDI-TIMS-MSI, while optimization of laser sampling minimized ion-load-dependent mobility shifts. Narrow mobility windows reduced reciprocal PG/BMP cross-talk to below 4% while preserving selective detection under strongly unbalanced abundance conditions. The workflow enabled distinct precursor- and product-ion imaging of endogenous PG 34:1 and BMP 34:1 in sagittal mouse brain, supporting their acyl-chain-level assignment as PG 16:0_18:1 and BMP 16:0_18:1. Application to a CLN3-knockout mouse model revealed BMP-specific reductions across brain, kidney, and lung that were not mirrored by the corresponding PG isomers, providing an orthogonal biological validation of the analytical discrimination. Mobility-constrained targeted MS/MS additionally resolved type-II isotopic interference that remained ambiguous at the MS1 level. Overall, this work provides a strategy for reciprocal spatial discrimination and structural confirmation of endogenous BMP and PG isomers directly in tissue and highlights the value of combining ion mobility with targeted product-ion imaging to increase molecular specificity in spatial lipidomics.

## Introduction

Spatial lipidomics by mass spectrometry imaging (MSI) has substantially advanced our understanding of the complex and heterogeneous lipid architecture of biological tissues^1,2^. Conventional MS-based lipidomics workflows generally require tissue homogenization and therefore lose the spatial context needed to investigate regio-specific lipid remodeling and microenvironment-dependent metabolic alterations ^3,4^. MSI overcomes this limitation by enabling label-free molecular mapping directly within tissue sections ^5^, which is particularly relevant given the central roles of lipids in membrane organization, cellular homeostasis, metabolism, and signaling ^6^. Alterations in lipid composition and metabolism are implicated in numerous pathological conditions, including Alzheimer’s disease ^7,8^, lysosomal storage disorders ^9^, and multiple cancer types ^10–12^. Lipid analysis by MSI is facilitated by the efficient ionization of several lipid classes using matrix-assisted laser desorption/ionization (MALDI) **^13^** and desorption electrospray ionization (DESI) ^14^. Nevertheless, confident on-tissue lipid characterization remains analytically challenging because of the structural complexity and wide dynamic range of the lipidome ^15^. Numerous isomeric, isobaric, and isotopically overlapping species are densely distributed within lipid-rich spectral regions, particularly between m/z 700 and 900. In this crowded mass range, low-abundance lipids are especially susceptible to ion suppression and interference from more abundant co-distributed species. Several advanced analytical strategies have been proposed to address these limitations. Ultra-high-resolution MSI is able to resolve lipid species separated by very small mass differences ^16–18^ whereas ozone-induced dissociation workflows provide structural information on carbon–carbon double-bond positional isomers within glycerophospholipid subclasses ^19,20^. In parallel, ion mobility (IMS) has emerged as a powerful orthogonal separation dimension **^21,22^**, improving isobaric and isomeric resolution in the gas phase ^23,24^. *In situ* MS/MS can be also used to enhance confidence for structural discrimination ^25,26^.

Despite their effectiveness, these approaches have predominantly focused on abundant lipid classes such as glycerophospholipids and sphingolipids. Numerous low-abundance lipid subclasses remain poorly covered because their signals are frequently masked by dominant species. Bis(monoacylglycero)phosphates (BMPs) represent an emblematic example. BMPs are negatively charged glycerophospholipids localized almost exclusively in late endosomal and lysosomal membranes and accounting for approximately 1% of total cellular phospholipids ^27,28^. Despite their relatively low abundance, BMPs are essential regulators of lysosomal membrane organization, lipid catabolism, and cholesterol homeostasis. Altered BMP metabolism has been associated with multiple lysosomal storage and neurodegenerative disorders, including neuronal ceroid lipofuscinoses ^29,30^. The analysis of BMPs remains particularly challenging because of their low abundance, and, additionally, structural isomerism with phosphatidylglycerols (PGs).

Based on accurate mass alone, BMPs and PGs cannot be distinguished by MS^1^ alone ^31^. Liquid chromatography–mass spectrometry (LC–MS) can overcome this limitation through chromatographic separation ^32^. Hydrophilic interaction liquid chromatography (HILIC) enables class-level separation based on polar headgroup chemistry and has been successfully applied to the discrimination and quantification of BMPs and PGs ^32,33^. In negative-electrospray (ESI) MS/MS, however, BMPs and PGs often generate highly similar and poorly class-diagnostic fragmentation patterns. Confident discrimination therefore generally relies on chromatographic retention, authentic standards, orthogonal ion-mobility information, or complementary positive-ion MS/MS fragmentation^33^. In MSI, the absence of chromatographic separation, together with the low abundance of BMPs and their isomerism with PGs, makes their confident spatial determination a formidable analytical challenge. Previous studies of the retinal pigment epithelium reported MALDI-FTICR-MS signals assigned to BMP species, but these assignments relied on MS^1^ accurate mass supported by LC–MS/MS analysis of tissue extracts, therefore did not directly discriminate BMPs from PGs in situ ^34^. The employment of Trapped Ion Mobility (TIMS) demonstrated that MALDI-TIMS can separate BMP and PG standard mixture, but this capability was not validated for endogenous BMP/PG pairs on tissue ^23^. Additionally, LC–TIMS-derived CCS information was transferred to MALDI-TIMS-MSI to support BMP annotation; however, although one BMP species was assigned, the corresponding PG was not detected, preventing direct visualization and reciprocal confirmation of the two isomers on tissue **^35^**. Thus, a clear simultaneous visualization and confident on-tissue discrimination of endogenous BMP and PG isomers have remained elusive.

To address this analytical gap, in the present work we developed a Trapped Ion mobility–guided MALDI-MSI strategy that integrates experimentally determined ion-mobility 1/K₀ information derived from orthogonal HILIC-DDA-PASEF experiments with mobility-constrained targeted MS/MS acquisition. In contrast to previous MS^1^-based workflows, the proposed approach directly detects both precursor and product ions of endogenous BMP/PG isomeric pairs, generating distinct precursor- and product-ion images and enabling acyl-chain-level spatial annotation. The method additionally resolves type-II isotopic interference from co-distributed lipid classes. As proof of concept the method was applied to map the spatial distribution of BMPs and PGs across brain, kidney, and lung tissues from wild-type and CLN3-knockout mice.

## Experimental section

### Reagents

LC–MS grade water (H_2_O), acetonitrile (ACN), methanol (MeOH), chloroform (CHCl_3_), Isopropanol (IPA), ethanol (EtOH), LC-MS grade additive ammonium acetate (CH_3_COONH_4_) were purchased from Merck (Milan, Italy). Authentic and deuterated BMPs and PGs standard (PG 18:1_18:1 cod. A84475, BMP 18:1_18:1 cod. A85075, PG 16:0_18:1 cod. A84457, BMP 14:0_14:0 (S,R) cod. A85073, BMP 18:1_18:1-D_5_ cod. A85067, PG 16:0-18:1-D_5_ cod. A86385) were purchased from Avanti Research (Alabaster, AL, USA). MALDI matrices 1,5-diaminonaphthalene (DAN), Norhamane (NRM), and naphthylethylenediamine dihydrochloride (NEDC) were purchased from Merck. MALDI IntelliSlides were purchased from Bruker Daltonics, ESI-L Low Concentration Tuning Mix was purchased from Agilent. Unless stated otherwise all other reagents were purchased from Merck.

### Biological Samples

C57BL/6J wild-type (WT) and CLN3 knockout (KO) mice (male, n=3 per group) were provided by Medina Lab. Housing and handling of mice were done in compliance with national guidelines. Kidney, lung, and brain samples were frozen in liquid nitrogen immediately after dissection, which took place within 5 min of sacrifice. All animal procedures were approved by the Ethical Committee of the Italian Ministry of Health (license number CE571.162). All efforts were made to minimize animal suffering and to reduce the number of animals used.

### Sample preparation for MSI analysis

The tissues were mounted in a cryostat microtome (Leica CM3050S, Leica Microsystems, Wetzlar, Germany) and attached to the holder using a minimal amount of water, without embedding medium, and sectioned at thicknesses of 12 μm at -20°C upon one-hour conditioning. Tissue sections were thaw-mounted onto pre-cooled IntelliSlides and stored at -80 °C until further use. Sections were vacuum-desiccated 1h prior to acquiring the optical images on an Epson Perfection V850 Pro scanner.

### Tissue washing and Matrix deposition

For washing studies, serial tissue sections were collected from the same tissue block. Ammonium acetate buffer was prepared to 50 mM concentration and chilled to 4 °C and the pH was measured^36^. Tissue sections were washed by immersing the edge of the slide containing the mounted section into chilled buffer and keeping it stationary for 5 s. The slide was then fully submerged in fresh buffer for an additional 15 s. After washing, excess liquid was gently blotted away, and the sample was placed back into the desiccator for 15 min.

DAN (5 mg/mL in 90% ACN) was sprayed over the tissue sections using an automated pneumatic sprayer (TM-Sprayer M5, HTX Technologies) to coat the samples with 2 µg/mm² of matrix. The sprayer was operated at a nozzle temperature of 40 °C, a nitrogen pressure of 10 psi, a flow rate of 100 μL/min, and a nozzle velocity of 1200 mm/min, twelve passes with a CC spray pattern and 2.5 mm track spacing were used. To investigate the effect of different MALDI matrices on the ionization efficiency of the target molecules, NRM and NEDC were also employed. The application conditions are described in **Table S1**.

### On-tissue standard spotting, selectivity and analytical sensitivity assessment

For on-tissue standard spotting experiments, PG 18:1_18:1 and BMP 18:1_18:1 standards were reconstituted in CHCl₃/MeOH (2:1, v/v), diluted in MeOH/H₂O (1:1, v/v), and 0.2 µL of each solution was spotted onto kidney tissue sections. The spots were allowed to dry completely before MALDI matrix application. For reciprocal signal contribution assessment, PG and BMP standards were individually spotted at 1.2 pmol/mm² and analyzed under identical experimental conditions. Reciprocal cross-talk was calculated as the signal intensity of each lipid detected in the non-target spot relative to that measured in the corresponding target spot. Selectivity was further evaluated using PG:BMP mixtures at molar ratios of 1:1, 10:1, 50:1, and 1:10. For analytical sensitivity assessment, serial dilutions of PG and BMP standards ranging from 0.1 to 20 pmol/mm² were spotted at six concentration levels, together with a MeOH/H₂O (1:1, v/v) control. Calibration curves were generated by plotting the measured signal response against the amount of analyte deposited per unit area (pmol/mm²), and the resulting models were used to determine the limits of detection (LOD) and quantification (LOQ) (**details in Table S2**).

### ESI-TIMS and MALDI-TIMS ion mobility agreement

For the comparison of ion mobility measurements obtained by ESI-TIMS and MALDI-TIMS, a panel of four PG and BMP standards, including PG 18:1_18:1, BMP 18:1_18:1, PG 16:0_18:1, and BMP 14:0_14:0, was analyzed at three concentration levels (1, 10, and 100 µM). For ESI-TIMS analysis, the standard solutions were directly infused into the mass spectrometer. For MALDI-TIMS analysis, each standard solution was mixed 1:1 (v/v) with the DAN matrix solution and spotted onto the MALDI target plate. The resulting ion mobility distributions and corresponding 1/K_0_ values were compared between the two ionization approaches.

To further evaluate the agreement between MALDI-TIMS and MALDI-TIMS-MSI measurements, the same PG and BMP standards (15 µM; 1.3 pmol/mm²) were spotted onto tissue sections prior to DAN matrix application. The resulting on-tissue standard spots were used both to compare the measured 1/K_0_ values with those obtained by MALDI-TIMS analysis of the corresponding standards and to optimize the number of laser shots for subsequent MALDI-TIMS-MSI experiments.

### Instrumentation and MALDI-TIMS-MS^1^ imaging

All MALDI-MSI experiments were performed on a timsTOF fleX with MALDI-2 and microGRID technology (Bruker Daltonics) equipped with a SmartBeam 3D 355 nm laser. MALDI-TIMS-MS^1^ image were acquired in negative ion mode from m/z 300−1350, the 1/K_0_ range was set to 0.7−1.80 V·s/cm^2^ with a ramp time of 250 ms. The laser size was set to 20 μm in timsControl (Bruker Daltonics), with 100 laser shots per position and 10kHz laser repetition rate, while the raster size was set to 40 μm in flexImaging (version 7.4) to allow for multiplexed data acquisition (collection of MS^1^ and MS^2^ spectra within the same MALDI MSI pixel). The ion transfer optics parameters were set as follows: Multiple RF amplitude 300 Vpp, RF collision cell voltage amplitude 1550 Vpp, focus pre TOF transfer time and pre-pulse storage were set to 85 and 10 µs, respectively. The quadrupole low mass was set to m/z 300. The MS method was externally calibrated using red phosphorus dissolved in 50% ACN and spotted beside the tissue section and internal calibration was applied using phosphatidylinositol PI (38:4) ion [M-H]^-^ as lock mass. Mobility calibration was performed using ESI with ESI-L Low Concentration Tuning Mix. The laser power was optimized at the start of each run and then held constant during the experiment. Tissue sections were analyzed in a random order to prevent any possible bias due to matrix degradation or variation in mass spectrometer sensitivity.The centroid spectra were imported into SCiLS Lab software (v.2024a, Bruker Daltonics). The feature list was created using the feature finding function using root mean square (RMS) intensity normalization followed by the T-ReX^3^ algorithm. The resulting mobility-filtered ion images were extracted, and molecular annotations were performed by MetaboScape 2025b (Bruker Daltonics, Bremen, Germany) directly from SCiLS and was based on a target list generated from hydrophilic interaction liquid chromatography (HILIC)-negative-DDA-PASEF acquisition of homogenized tissues. Statistical analysis and bar graphs were performed using GraphPad Prism 8.0 (GraphPad Software, Boston, MA, USA, www.graphpad.com).

### MALDI iPRM-PASEF

MALDI iPRM-PASEF data were acquired in negative ion mode with the scan mode set to “prm-PASEF” from m/z 50−1350, and 1/K_0_ range 0.5−1.80 V·s/cm^2^ with a ramp time of 250 ms. The ion transfer optics parameters were set as follows: Multiple RF amplitude 300 Vpp, RF collision cell voltage amplitude 800 Vpp, focus pre-TOF transfer time and pre-pulse storage were set to 65 and 5 µs, respectively. The quadrupole low mass was set to m/z 100. Precursors were isolated using a 2 m/z isolation window and collision energies for MS^2^ fragmentation were interpolated based on mobility isolation, ranging from 25 eV at 0.8 V·s/cm^2^ to 60 eV at 1.80 V·s/cm^2^.

Multiplexed MALDI-MSI analysis was then performed by generating a new imaging run file derived from the flexImaging sequence of the MALDI-TIMS-MS^1^ run. The laser offset was set to X = 0, Y = 0 μm for MALDI-TIMS-MS^1^ run and to X = 20, Y = 0 μm for MS^2^ analysis. This offset enabled the acquisition of MALDI-tims-MS^2^ spectra from previously unablated sample regions, allowing the complete MALDI iPRM-PASEF workflow to be performed on a single sample while preserving the same spatial distribution as in the MALDI-TIMS-MS^1^ dataset.

### Histological Staining and Co-Registration with MSI Data

Following MSI analysis, the MALDI matrix was removed by submerging the slides in 95% ethanol for 30 s. The tissue sections were subsequently fixed in PFA 4% and rehydrated prior to histological staining. Kidney and lung sections were stained with hematoxylin and eosin (H&E), whereas brain sections were subjected to Nissl staining. The stained slides were cover-slipped and scanned using an Epson Perfection V850 Pro scanner. Whole-slide histological images were subsequently imported into SCiLS Lab and co-registered with the corresponding MSI datasets to enable spatial correlation between molecular distributions and tissue morphology.

### Sample preparation for HILIC-TIMS analysis

Lipids were extracted from sections of kidney, brain and lung tissues, pooled separately and extracted using a CHCl₃/MeOH based extraction. Briefly, 330 µL of ice-cold MeOH containing 10 ng of BMP 18:1_18:1-D_5_, BMP 14:0_14:0 (S,R) and PG 16:0-18:1-D_5_ were added to samples and then sonicated in an ultrasound bath for 10 min. Subsequently, 670 µL of ice-cold CHCl_3_ were transferred to the tube and the solution was continuously agitated in a thermomixer (Eppendorf, Milan, Italy) for 70 min, 1000 rpm at 4°C. Then, 188 µL of H_2_O were added to induce phase separation and, after 10 min of incubation, the lipid extracts were centrifuged. The bottom layer was collected and dried using a SpeedVac (Savant, Thermo Scientific, Milan, Italy). The dried extracts were dissolved in 100 µL of IPA/ACN/H_2_O (13:6:1 v/v) before the UHPLC-TIMS-MS analysis.

### HILIC -TIMS method parameters

HILIC-TIMS-MS analyses were performed on an Elute^+^ UHPLC system (Bruker Daltonics) coupled online to the same timsTOF fleX ESI/MALDI instrument (Bruker Daltonics) equipped with an Apollo II electrospray ionization (ESI) probe. The separation was performed with an Ascentis Express® HILIC column (150 × 2.1 mm, 2.7 μm; Column Length × Internal Diameter, particle size). The column temperature was set at 35°C, a flow rate of 0.3 mL/min was used, mobile phase consisted of (A): 35 mM CH_3_COONH_4_ in H_2_O, pH 6.8 (B): ACN. The following gradient has been used: 0 min, 97% B; 0.5 min, 97% B; 20 min, 75% B; 20.6 min, 60% B; 20.60 min, 40% B; 22.60 min, 40% B; 22.70 min, 97% B; and then 3 min for column re-equilibration.

The TIMS-MS analyses were performed in data-dependent parallel accumulation serial fragmentation (DDA-PASEF) in negative ionization, and each sample was injected in triplicate. The injection volume was set at 5 µL. Source parameters: Nebulizer gas (N_2_) pressure: 4.0 Bar, Dry gas (N_2_): 10 L/min, Dry temperature: 220°C. Mass spectra were recorded in the range m/z 50–1500, with an accumulation and ramp time to 100 ms each. The ion mobility was scanned from 0.55 to 1.80 Vs/cm^2^. Precursors for data-dependent acquisition were isolated using a 2 m/z isolation window and fragmented with a fixed collision energy (50 eV). The total acquisition cycle was of 0.32 s and comprised one full TIMS-MS scan and two PASEF MS/MS scans. Exclusion time was set to 0.1 min, Ion charge control (ICC) was set to 7.5 Mio. The instrument was calibrated for both mass and mobility using the ESI-L Low Concentration Tuning Mix with the following composition: m/z, 1/K_0_: (301.99814, 0.6678 V·s/cm^2^), (601.97897, 0.8781 V·s/cm^2^), (1033.98811, 1.2525 V·s/cm^2^), (1333.96894, 1.4015 V·s/cm^2^).

### HILIC-TIMS data analysis and processing

4D data alignment, filtering and annotation were performed with MetaboScape 2025b (Bruker) employing a feature finding algorithm (T-Rex 4D) that automatically extracts buckets from raw files. At the beginning of each LC-MS run, a mixture (1:1 v/v) of 10 mM sodium acetate calibrant solution and ESI-L Low Concentration Tuning Mix was injected to recalibrate, respectively, the mass and mobility data. Feature detection was set to 100 counts. The minimum number of data points in the 4D-TIMS space was set to 100, and recursive feature extraction was used (75 points).

### Lipid annotation

Lipid annotation was performed first with a rule-based annotation, based on characteristic fragments and their intensity in acquired MS/MS spectra, and, subsequently, using the LipidBlast spectral library of MS DIAL (http://prime.psc.riken.jp/compms/msdial/main.html) with the following parameters: Mass accuracy: narrow 2 ppm, wide 10 ppm; mSigma: narrow 30, wide 250, MS/MS score: narrow 800, wide 150. Collision cross-section (CCS)%: narrow 2, wide 3.5. The spectra were processed in negative mode using [M-H]^−^, [M+Cl]^−^ and [M-H_2_O]^−^ as adducts. CCS values were compared with those predicted by CCSbase platform (https://ccsbase.net/), the assignment of the molecular formula was performed for the detected features using Smart Formula™ (SF). Each lipid feature was manually curated following Lipidomics Standard Initiative (LSI) guidelines (https://lipidomics-standards-initiative.org/guidelines/lipid-species-identification/general-rules) as reported previously ^37^.

## Results and Discussion

### Workflow overview

The spatial lipidomics workflow is outlined in **Figure 1** and consists of four main phases. 1) A MALDI-TIMS-MS^1^ survey scan analysis to assess the presence of target PG and BMP species based on their theoretical m/z values and corresponding ion mobility (1/K₀) profiles. 2) An orthogonal HILIC-DDA-PASEF analysis in negative ESI mode is performed to obtain accurate MS, isotopic pattern, MS/MS fragmentation and ion mobility (1/K_0_) information for feature annotation and to characterize the mobility-based separation of PG and BMP species. 3) The resulting annotated feature list, including accurate m/z and 1/K₀ values, is used to build the targeted iPRM method, incorporating mobility-based precursor selection to improve PG/BMP discrimination and minimize cross-talk. 4) Finally, MALDI-iPRM analysis is performed directly on tissue sections using mobility-constrained precursor selection and targeted MS/MS acquisition for spatially resolved lipid identification.

**Figure 1.**
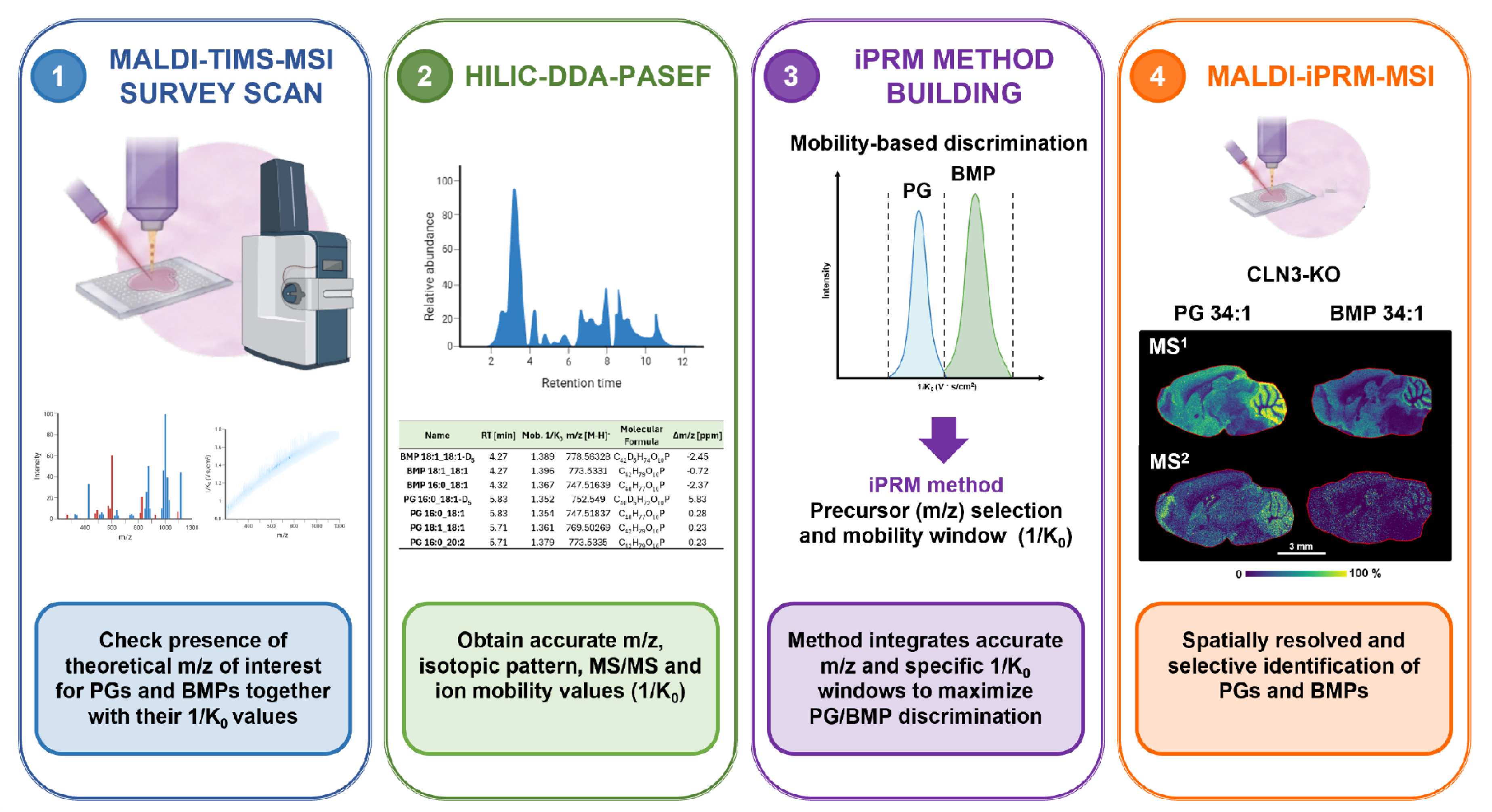
Integrated workflow for mobility-assisted selective spatial lipidomics of PGs and BMPs. The workflow combines MALDI-TIMS-MS^1^ survey analysis with orthogonal HILIC-DDA-PASEF characterization to obtain accurate mass, MS/MS, and ion mobility information. These data are used to establish mobility-based PG/BMP discrimination and to develop a mobility-constrained iPRM method for targeted MALDI-iPRM analysis and spatially resolved lipid identification. *Created in bioRender.com*

### Preliminary optimization of MALDI-TIMS-MSI acquisition conditions

For initial method development and optimization, fresh frozen mouse kidney samples were used and were cryosectioned longitudinally. Considering that BMPs and PGs preferentially ionize in negative mode, we first evaluated the effect of different matrices on their ionization efficiency. To identify the suitable conditions for the detection of PG and BMP species, three matrices, NRM, NEDC and DAN, were compared. The performance of each matrix was evaluated by comparing the number and intensity of putatively annotated PG species detected under identical acquisition conditions. For representative PG species, DAN provided the highest signal intensity for the species at m/z 747.51, putatively annotated as PG 34:1, compared with both NRM and NEDC. Moreover, NRM showed a lower signal response for PG 36:2 at m/z 773.53. The species at m/z 721.50, putatively annotated as PG 32:0, was exclusively detected using DAN (**Figure S1**). Based on the overall performance for PG detection, DAN was selected for subsequent experiments. Subsequently, using the selected matrix, the sample preparation procedure was optimized by assessing the effect of an ammonium acetate washing step prior to matrix application. Ammonium-containing buffers have previously been reported to enhance sensitivity in negative ion mode lipid imaging **^36^**. Thus, we evaluated whether a 50 mM CH_3_COONH_4_ washing step prior to matrix application could further improve the ionization efficiency of PG/BMP species. As shown in **Figure S2a, b**, the washing step resulted in an overall increase in total ion current, corresponding to a 1.76 ± 0.57 -fold increase compared with the untreated tissue sections. Notably, putatively annotated PG species showed an approximately 2.63 ± 1.25 -fold increase in signal intensity under washing conditions. Given the closely related chemical properties and ionization behavior of PGs and BMPs, this improvement was considered potentially applicable to BMP detection as well.

### 3.3 Cross-platform agreement of ESI-TIMS and MALDI-TIMS ion mobility

To assess the agreement of ion mobility measurements obtained using the two ionization approaches, a panel of four PG and BMP standards was analyzed at three concentration levels (1, 10, and 100 µM) using both ESI-TIMS and MALDI-TIMS. As shown in the correlation plots (**Figure S3a-c**), the 1/K_0_ values obtained by Direct Infusion (DI)-ESI-TIMS and MALDI-TIMS exhibited excellent agreement across all three concentration levels. Pearson’s correlation coefficients (*r*) were 0.9995, 0.9997 and 0.9999 at 1, 10, and 100 µM, respectively, while the corresponding coefficients of determination (R^2^) were 0.9990, 0.9994 and 0.9999. Notably, the R^2^ values remained ≥ 0.999 across the entire concentration range, indicating that the excellent agreement between ESI-TIMS and MALDI-TIMS was maintained across the concentration range, without substantial loss of correspondence between the two platforms. These results demonstrate highly consistent mobility measurements and support the transferability of 1/K_0_ information from ESI-TIMS to MALDI-TIMS analysis.

### MALDI-TIMS and MALDI-TIMS-MSI mobility agreement and laser shots optimization

To further assess the transferability of ion mobility information to tissue-based MALDI-MSI measurements, the 1/K_0_ values obtained from MALDI-TIMS analysis of spotted standards were compared with those measured directly during MALDI-TIMS-MSI experiments on tissue sections. The ion mobility values (1/K_0_) obtained under the two experimental configurations showed strong agreement as demonstrated by the correlation plot (**Figure S3d**), with both *r* and R^2^=0.9999, supporting the reproducibility of the measured 1/K_0_ values when moving from standard-based MALDI-TIMS measurements to tissue analysis.

Despite the good agreement between MALDI-TIMS and MALDI-TIMS-MSI measurements, the ion mobility distributions measured during tissue imaging may be influenced by acquisition parameters that affect the ion population introduced into the TIMS analyzer. In particular, increasing the number of laser shots increases the number of ions generated from the tissue and, consequently, the ion load entering the TIMS cartridge. This may alter the measured mobility distribution, potentially through ion-ion interactions and space-charge effects, thereby affecting the reproducibility of predefined mobility windows.

To investigate this effect, ion mobility values 1/K_0_ obtained from standards spotted on the MALDI target plate were compared with those measured from the same standards deposited on tissue sections under identical acquisition conditions. Representative extracted ion mobilograms (EIMs) of PG 18:1_18:1 and BMP 18:1_18:1 are shown in **Figures S4a** and **S4b**, respectively. At 200 laser shots, standards analyzed on tissue showed a clear shift in ion mobility distributions compared with those measured on the MALDI target plate. Conversely, no appreciable differences in 1/K_0_ values were observed when 100 laser shots were applied.

Consistently, **Figure S5** shows that increasing the number of laser shots from 50 to 100 preserved the measured mobility values, whereas acquisition with 200 laser shots resulted in a marked displacement of the ion mobility distributions.

The effect of laser shots was further evaluated during MALDI-TIMS-MSI acquisition by comparing the ion mobility distribution of the endogenous PG 16:0_18:1 species detected in tissue with that of the corresponding pure standard spotted on the tissue section. As shown in **Figure S6**, the endogenous and standard PG 16:0_18:1 species displayed comparable mobility profiles when acquired using 50 and 100 laser shots, indicating preservation of the characteristic 1/K_0_ distribution under these conditions. Conversely, acquisition with 200 laser shots resulted in a shift toward higher 1/K₀ values, confirming that excessive ion populations generated during tissue imaging can affect the measured ion mobility profiles. This observation is particularly relevant for PG/BMP discrimination, as BMP species generally exhibit higher 1/K_0_ values than their corresponding PG isomers. Consequently, a shift toward higher 1/K₀ values may reduce the effective separation between the two lipid classes and compromise the selectivity of predefined mobility windows.

Based on these results, 100 laser shots were selected as the optimal acquisition condition, providing sufficient signal intensity while preserving the characteristic ion mobility distributions required for reliable mobility-based PG/BMP discrimination. These parameters were subsequently applied to all MALDI-TIMS-MSI and MALDI iPRM-PASEF experiments.

### 3.5 Orthogonal HILIC-DDA-PASEF acquisition and feature annotation

Parallel HILIC-DDA-PASEF experiments were performed on the same TIMS-MS platform to support the molecular characterization and annotation of lipid species detected by MALDI-TIMS-MSI. As shown by the extracted ion chromatograms (EICs) obtained from both the standard mixture and the kidney tissue extract (**Figure S7**), HILIC chromatography is well suited for the analysis of BMPs and PGs, as previously reported ^33^. In HILIC separations, these lipid classes are resolved according to differences in the polarity and chemical properties of their head groups, enabling their chromatographic discrimination and facilitating the assignment of lipid classes. The observed retention behavior of the endogenous lipid species was consistent with that of the corresponding standards, supporting confident annotation. Under the HILIC separations, BMP species consistently eluted between 3.5 and 4.8 min, whereas PG species were detected within a later retention window of 5.3-6.3 min, providing an additional orthogonal level of separation prior to TIMS analysis.

HILIC-DDA-PASEF analysis provided complementary molecular information, including accurate mass, isotopic pattern, MS/MS fragmentation, and ion mobility values (1/K_0_). The features detected and annotated from the HILIC-DDA-PASEF dataset were subsequently compiled into an analyte list containing the corresponding accurate m/z and 1/K_0_ values. Complete annotation is reported in **Table S3**. This information was used as a reference for the annotation of the MALDI-TIMS imaging data, enabling the integration of molecular and ion mobility information obtained from the orthogonal HILIC-DDA-PASEF analysis into the spatially resolved MALDI-TIMS dataset.

### Ion Mobility-based discrimination of PGs and BMPs and Reciprocal cross-talk assessment

PGs and BMPs are isobaric lipids with highly similar MS/MS fragmentation patterns, requiring an additional dimension of separation for confident discrimination **^38^**. To evaluate the capability of MALDI-TIMS-MSI to discriminate PG and BMP species directly on tissue, individual PG 18:1_18:1 and BMP 18:1_18:1 standards, as well as their equimolar mixture (PG:BMP, 1:1), were deposited onto tissue sections and analyzed under the optimized acquisition conditions. As shown in **Figure 2a, b**, MSI images generated using the characteristic ion mobility windows of PG 18:1_18:1 and BMP 18:1_18:1 selectively visualized the corresponding lipid species in the individual standard spots. When the PG/BMP mixture was analyzed, both lipid classes could be independently visualized by applying their respective mobility (1/K_0_) extraction windows, resulting in spatial localization patterns consistent with the individual standards. To further characterize the extent of mobility separation achieved, EIMs of the PG/BMP 18:1_18:1 mixture were evaluated (**Figure 2c**). Although the two lipid species displayed distinct mobility (1/K_0_) values, their mobility distributions partially overlapped, indicating that complete baseline separation was not achieved. The separation appeared more evident in the MALDI-TIMS-MS analysis of the standard mixture (orange trace), whereas a similar but slightly broader overlap was observed in the corresponding MALDI-TIMS-MSI on-tissue analysis (blue trace), likely reflecting the more complex ion generation conditions. This partial mobility overlap highlighted the need to define optimized extraction mobility windows to maximize PG/BMP selectivity while minimizing reciprocal cross-talk. Reciprocal cross-talk refers to the residual contribution of one lipid species detected within the mobility window assigned to the other species, resulting from the partial overlap of their ion mobility distributions. Reciprocal cross-talk was calculated by comparing the signal intensity of each lipid species detected in the non-target standard spot with that measured in the corresponding target standard spot, using progressively narrower extraction mobility windows to improve selectivity and reduce signal contamination between PG and BMP channels (±0.008, ±0.006, and ±0.004 1/K₀), according to the following formulas:

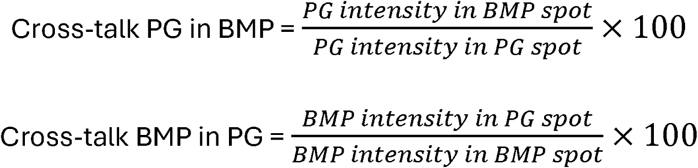

**Figure 2.**
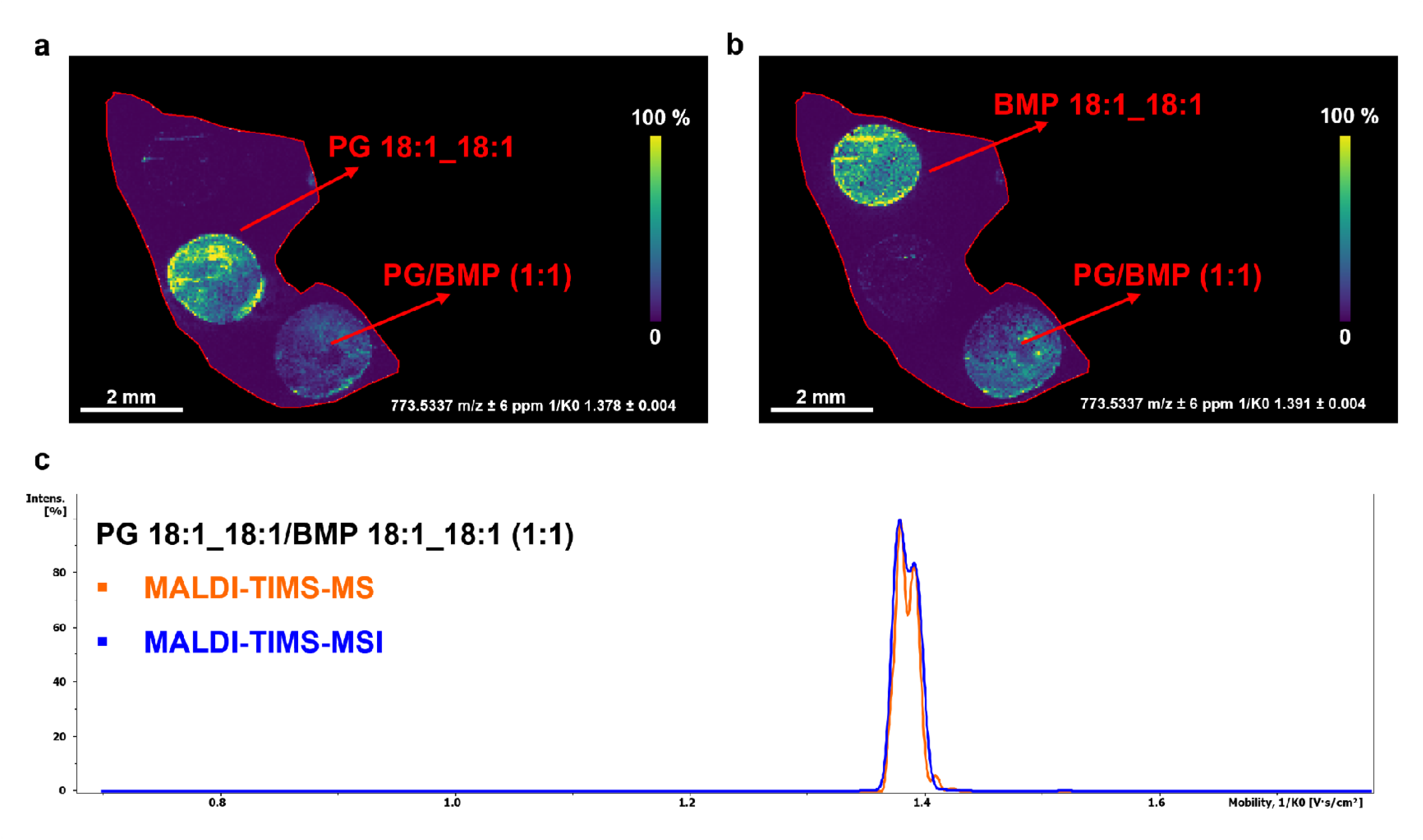
Ion mobility-based discrimination of PG and BMP isobaric lipid species by MALDI-TIMS-MSI. a, b) MALDI-TIMS-MSI images of individual PG 18:1_18:1 and BMP 18:1_18:1 standards and of the corresponding equimolar mixture (PG/BMP, 1:1) spotted onto tissue sections. Images were generated by extracting the signals within the characteristic ion mobility values 1/K_0_ ± 0.004 of PG and BMP, allowing selective visualization of the two lipid species. In the mixed standard spot, PG and BMP distributions were independently visualized by applying their respective mobility extraction windows. Scale bar, 1 mm. Color scale bar is shown as percentage of maximum intensity. Pixel size, 20 μm. c) Extracted ion mobilograms (EIMs) of the PG/BMP 18:1_18:1 standard mixture acquired by MALDI-TIMS-MS (**orange trace**) and MALDI-TIMS-MSI (**blue trace**).

As summarized in **Table 1**, reducing the mobility window markedly decreased reciprocal signal contribution between the two lipid classes. Cross-talk decreased from 15.83% and 14.26% (PG in BMP and BMP in PG, respectively) using a ±0.008 window to 9.26% and 9.37% with a ±0.006 window. Using the narrowest mobility extraction window ±0.004, only limited reciprocal signal contribution was observed, corresponding to 3.55% for PG detected in the BMP spot and 2.65% for BMP detected in the PG spot, demonstrating the high selectivity of the mobility-based workflow.

**Table 1.** Effect of ion mobility extraction window on reciprocal cross-talk between PG 18:1_18:1 and BMP 18:1_18:1 standards.

| Mobility extraction window ($\pm 1/K_0$) | PG detected in BMP (%) | BMP detected in PG (%) |
| --- | --- | --- |
| $\pm 0.008$ | 15.83 | 14.26 |
| $\pm 0.006$ | 9.26 | 9.37 |
| $\pm 0.004$ | 3.55 | 2.65 |

### Selectivity across different PG/BMP molar ratios

Although reciprocal cross-talk provides a measure of analytical selectivity under idealized conditions, biological samples frequently contain PG 18:1_18:1 and BMP 18:1_18:1 species at markedly different relative abundances Therefore, the robustness of the proposed mobility-based discrimination strategy was further evaluated using mixtures of authentic PG and BMP standards prepared at different molar ratios (PG/BMP 1:1, 10:1, 50:1 and BMP/PG 10:1).

For each mixture, signal intensities were extracted using the optimized mobility windows assigned to PG and BMP, and the measured PG/BMP signal ratios were compared with the corresponding theoretical molar ratios. Representative MALDI-TIMS-MSI images are shown in **Figure S8**, while the measured signal ratios are summarized in **Table S4**. The measured PG/BMP ratios closely followed the expected trend across the mixtures investigated. Measured ratios of 1.12 and 10.44 were obtained for theoretical PG:BMP ratios of 1:1 and 10:1, respectively. At the highest abundance contrast (PG:BMP 50:1), the measured ratio decreased to 31.57, whereas a theoretical BMP:PG ratio of 10:1 yielded a measured ratio of 8.92. Although deviations from the theoretical values became apparent at the highest abundance contrast, selective detection of the minor component was maintained in all cases, demonstrating the selectivity of the proposed mobility-based workflow over a broad range of relative analyte abundances.

### Mobility-constrained MALDI-iPRM method development

Following optimization of the mobility windows, a targeted mobility-constrained MALDI-iPRM-PASEF method was developed to translate PG/BMP mobility discrimination into selective on-tissue MS/MS acquisition. The inclusion list was generated from the orthogonal HILIC-DDA-PASEF dataset and incorporated accurate precursor m/z, experimentally determined 1/K₀ values, and characteristic product ions for each annotated lipid species. PG and BMP targets were combined within a single scheduled acquisition, allowing isomeric precursors sharing identical m/z values to be selectively isolated and fragmented according to their distinct mobility coordinates. During MALDI-iPRM acquisition, precursor selection is constrained by both accurate mass and ion mobility, thereby restricting MS/MS acquisition to ions falling within predefined mobility windows. This strategy minimizes precursor interference arising from the partial overlap of the PG and BMP mobility distributions and enables selective fragmentation of each lipid species directly in tissue

The performance of this strategy was initially evaluated using PG 18:1_18:1 and BMP 18:1_18:1 standards spotted onto tissue sections (1.3 pmol/mm²). Despite their identical precursor m/z, the two species were selectively targeted at their respective mobility coordinates (**Figure 3a-d**). In both cases, the product ion at m/z 281.2491, corresponding to the deprotonated 18:1 fatty acyl chain (**Figure 3b,d**), reproduced the spatial distribution of the respective mobility-selected precursor at m/z 773.5337. Although this fragment is shared by both isomers and is therefore not class-diagnostic, its spatial concordance with the selected precursor population confirmed the expected acyl-chain composition. Thus, the combination of accurate mass and ion mobility precursor selection with product ion imaging highlights that the proposed mobility-constrained MALDI-iPRM strategy enables selective targeting of isomeric PG and BMP species and provides an additional level of molecular confirmation prior to biological application.

**Figure 3.**
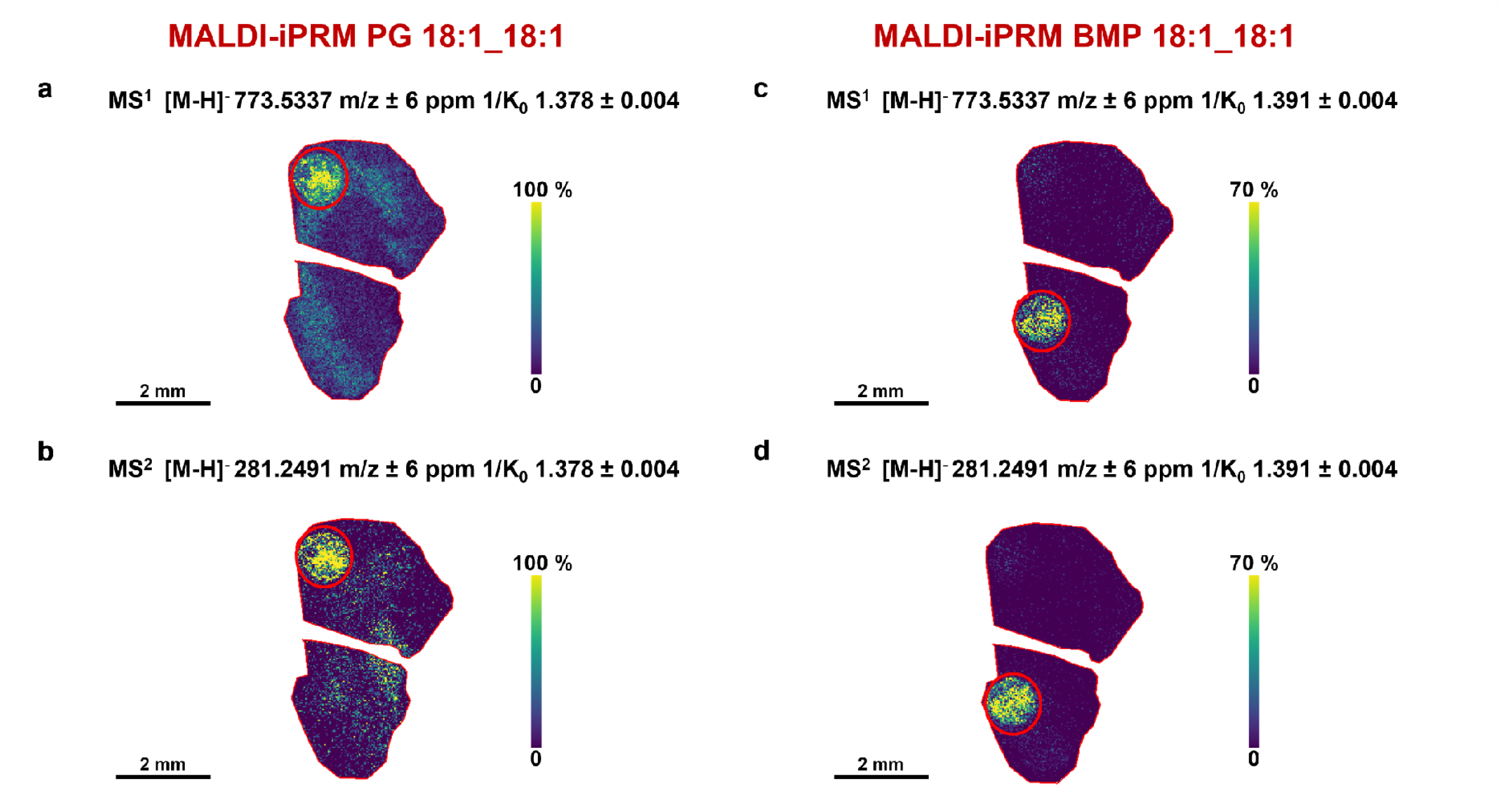
Validation of the MALDI-iPRM approach for the discrimination of the isobaric lipid pair PG and BMP. a) Precursor ion image of the PG 18:1_18:1 standard spot at m/z 773.5337 selected by MALDI-iPRM (1.378 ± 0.004 1/K_0_). b) Corresponding MS/MS product ion image showing the distribution of the 18:1 fatty acyl chain fragment at m/z 281.2491. c) Precursor ion image of the BMP 18:1_18:1 standard spot selected by MALDI-iPRM (1.391 ± 0.004 1/K_0_). d) Corresponding MS/MS product ion image at m/z 281.2491. Scale bar, 2 mm. Color scale bar is shown as percentage of maximum intensity. Pixel size, 20 μm; raster size, 40 μm. Data were normalized by RMS.

An additional advantage of the developed approach is demonstrated for the annotation of PG 36:2 in the kidney tissue sections, where the precursor signal was affected by a type-II isotopic interference arising from a co-distributed lipid species. MALDI-TIMS-MS analysis detected the ion at m/z 773.5337 (**Figure S9a**), assigned to PG 36:2, which partially overlapped with the M+1 isotopologue contribution at m/z 773.5309 (**Figure S9b**), of the ion detected at m/z 772.5291 (**Figure S9c**). The presence of this interference was supported by the isotopic pattern analysis shown in **Figure S9d** for both MALDI-MSI (**top**) and HILIC-DDA-PASEF spectra (**bottom**). The interfering precursor at m/z 772.5291 was annotated as PE O-40:8 based on its retention time (10.3 min), which was distinct from the elution window of BMPs and PGs, and its corresponding HILIC-DDA-PASEF fragmentation pattern (**Figure S9e**). Consequently, MS^1^-based annotation, even when combined with ion mobility information, was insufficient to unambiguously discriminate the PG 36:2 signal from the overlapping isotopic contribution. Mobility-constrained MALDI-iPRM followed by narrow post-acquisition extraction in m/z (±6 ppm) and ion mobility (±0.004 1/K₀) selectively recovered the PG 36:2-associated precursor and product-ion signals while excluding the isotopic contribution from PE O-40:8 (**Figure S9f**). The resulting MALDI-iPRM spectrum showed the characteristic product ions of PG species, confirming the structural assignment of PG 16:0_20:2, in agreement with the annotation obtained from HILIC-DDA-PASEF analysis (**Figure S9g**). Conversely, BMP 36:2 did not show evidence of type-II isotopic interference in the analyzed kidney tissue sections. The precursor displayed a distinct ion mobility profile and was confidently annotated as BMP 18:1_18:1 through MALDI-iPRM fragmentation (**Figure S9h**) and supported by the corresponding HILIC-DDA-PASEF fragmentation spectrum (**Figure S9i**). This example demonstrates that the benefit of the mobility-constrained iPRM workflow extends beyond BMP/PG isomerism, providing an additional level of selectivity against isotopic interferences that remain unresolved in conventional MS^1^ imaging.

### Proof-of-concept: on-tissue discrimination of endogenous PG 34:1 and BMP 34:1 in CLN3-knockout mouse brain

To evaluate the proposed workflow under biologically relevant conditions, we employed a CLN3-KO mouse model of juvenile neuronal ceroid lipofuscinosis (CLN3 disease, MIM #204200), in which previous lipidomic studies have reported a marked depletion of BMP species ^39^. We therefore investigated whether the method could selectively visualize the isomeric PG 34:1 and BMP 34:1 species directly in sagittal brain sections from wild-type and CLN3-KO mice (**Figure 4a, b**).

**Figure 4.**
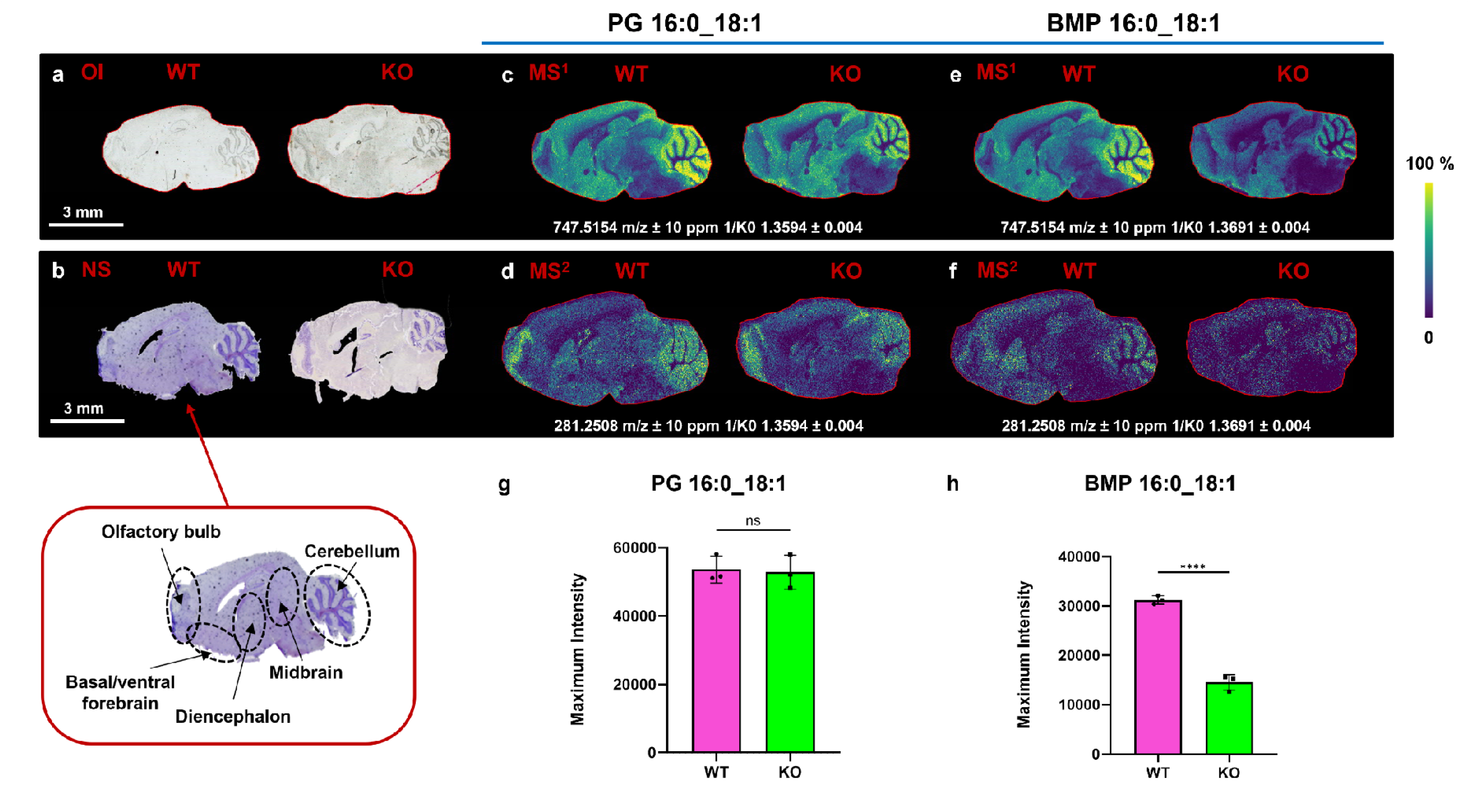
MALDI-iPRM discrimination of PG 16:0_18:1 and BMP 16:0_18:1 in sagittal brain sections from WT and CLN3-knockout (KO) mice. Optical image (a) and corresponding Nissl-stained section with an enlarged view showing the anatomical annotations (b). MALDI-TIMS MS^1^ ion image of **PG 16:0_18:1** (c) and the corresponding MALDI-iPRM product ion image at m/z 281.2508 (d). MALDI-TIMS MS^1^ ion image of **BMP 16:0_18:1** (e) and the corresponding MALDI-iPRM product ion image at m/z 281.2508 (f). All images are shown for both WT and CLN3-KO brain sections. Scale bar, 3 mm. Color scale bar is shown as percentage of maximum intensity. Pixel size, 20 μm; raster size, 40 μm. Bar graphs showing the signal intensities of **PG 16:0_18:1** (g) and **BMP 16:0_18:1** (h) in WT and CLN3-KO mice. Error bars represent the standard deviation of three biological replicates. Statistical analysis was performed using an unpaired two-tailed Student’s t-test (****P ≤ 0.0001, ns, not significant). Data were normalized by RMS. Abbreviations: OI, Optical Image; NS, Nissl Staining.

PG 34:1 and BMP 34:1 share the same elemental composition and precursor *m/z* (*m/z* 747.5154) but displayed distinct ion mobility values. Using mobility-constrained windows centered at 1/K₀ 1.3594 ± 0.004 and 1.3691 ± 0.004, respectively, the PG- and BMP-associated precursor populations could be independently visualized on tissue (**Figure 4c, e**).

Targeted on-tissue MS/MS of both mobility-selected precursor populations generated highly similar fragmentation patterns, consistent with the known negative-ion fragmentation behavior of PGs and BMPs. Both species produced abundant product ions at *m/z* 281.2508 (**Figure 4d, f**) and 255.2344 (**Figure S10a, b**), corresponding to the deprotonated fatty acyl anions of 18:1 and 16:0, respectively, together with the lower-abundance phosphate-containing product ion at *m/z* 152.9956 (**Figure S10a, b**). Detection of the 16:0 and 18:1 fatty acyl anions supported the acyl-chain-level annotation of both species as PG 16:0_18:1 and BMP 16:0_18:1. Importantly, the occurrence of the same characteristic product ions for both isomers further illustrates that negative-ion MS/MS alone does not provide sufficient class specificity and that PG/BMP discrimination relies on mobility-selective precursor isolation.

Despite their similar fragmentation behavior, PG and BMP displayed distinct spatial distributions throughout the sagittal brain section, which were preserved in the corresponding product-ion maps (**Figure 4c-f**). The spatial concordance between precursor and product-ion images provided direct on-tissue confirmation that the selected product ions originated from the respective mobility-defined precursor populations. At the regional level, PG 16:0_18:1 retained a largely conserved distribution between WT and CLN3-KO brains, including prominent cerebellar and forebrain signals. In contrast, BMP 16:0_18:1 depletion was spatially heterogeneous, with a more pronounced reduction in diencephalic and midbrain regions, whereas residual signal was comparatively better preserved in the cerebellum. A similar regional pattern was observed in the corresponding product-ion images.

At the whole-section level, comparison of wild-type and CLN3-KO brains revealed a marked decrease in BMP 16:0_18:1 signal intensity in the KO group, consistently observed in both precursor (**Figure 4e**) and product-ion images (**Figure 4f**), whereas the corresponding PG 16:0_18:1 isomer remained substantially unchanged (**Figure 4c,d**). The trend was also reflected in the signal intensity profiles shown in the bar graphs (**Figure 4g, h**), confirming a significant reduction of BMP 16:0_18:1 in CLN3-KO brain (**Figure 4h**), with no significant difference in PG 16:0_18:1 between genotypes (**Figure 4g**). A similar pattern was observed for PG/BMP 38:4 (**Figure S10c**). The fragmentation pattern supported its annotation as 18:0_20:4, and the distinct reduction of BMP 18:0_20:4 in the KO group (**Figure S10c, d**). The distinct response of the two mobility-resolved isomers to the CLN3 model provides orthogonal biological support for their assignment as distinct endogenous molecular species.

### Extending reciprocal PG/BMP discrimination and annotation across tissues

Having established on-tissue discrimination of different PG and BMP in brain, we next evaluated whether the workflow could be transferred to tissues with markedly different lipid composition and morphology. Kidney and lung sections were therefore analyzed using the same strategy. In kidney, endogenous PG/BMP discrimination was reproduced for both the 34:1 and 36:2 compositions (**Figure 5a**). Complete MALDI-iPRM product-ion images are reported in **Figure S11**. PG 16:0_18:1 and PG 16:0_20:2 showed no significant differences between WT and CLN3-KO mice, whereas BMP-associated signals were reduced in the KO group, reaching statistical significance for BMP 18:1_18:1 (**Figure 5b**).

**Figure 5.**
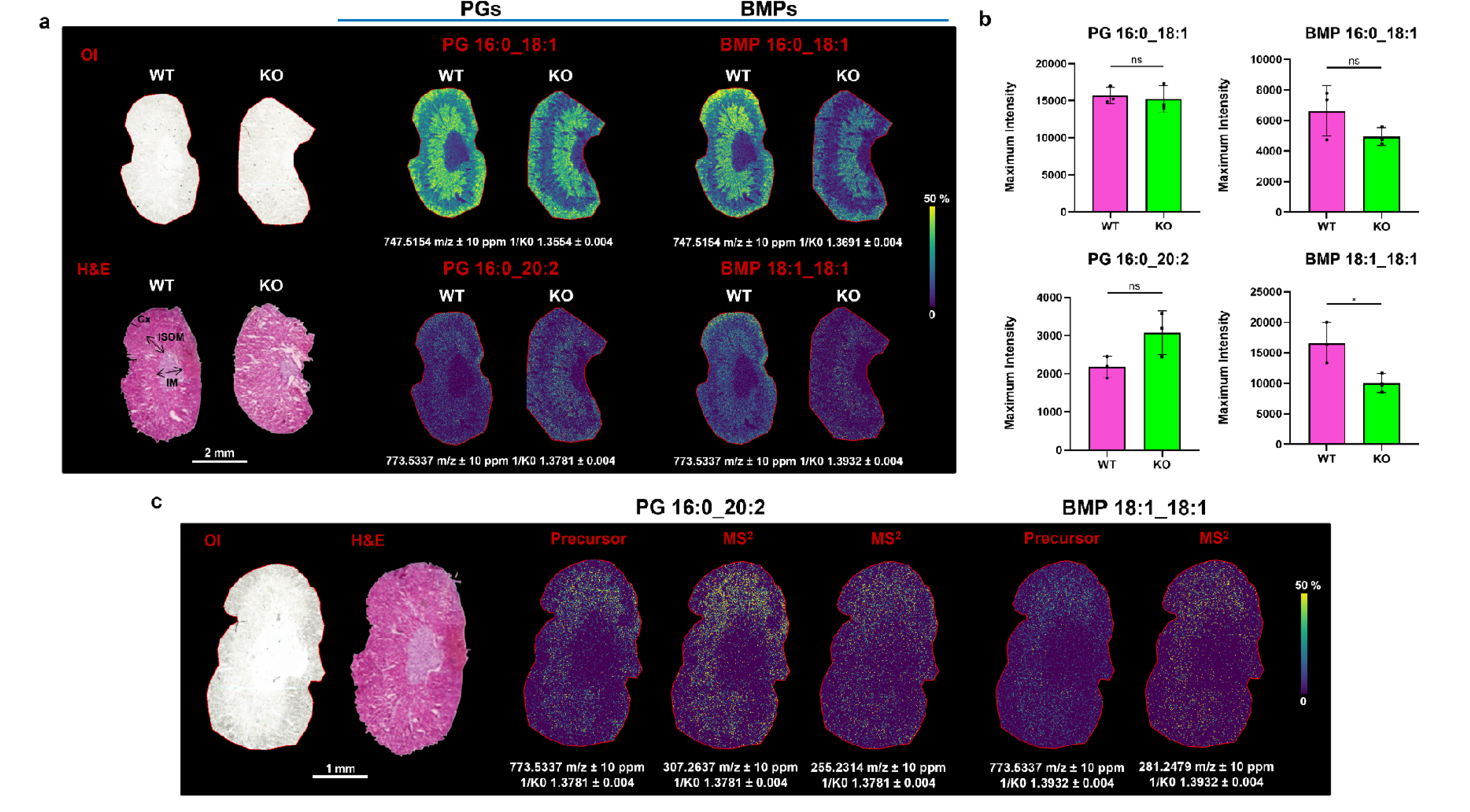
MALDI-iPRM discrimination of PGs and BMPs in kidney sections from WT and CLN3-knockout (KO) mice. a) MALDI-TIMS MS^1^ ion images of PG 16:0_18:1, PG 16:0_20:2, BMP 16:0_18:1 and BMP 18:1_18:1 in kidney sections from WT and CLN3-KO mice. Scale bar, 2 mm. Color scale bar is shown as percentage of maximum intensity. Abbreviations: OI, Optical Image; H&E, Hematoxylin and Eosin Staining with the anatomical annotations: Cx, renal cortex; ISOM, inner stripe of the outer medulla; IM, inner medulla. b) Bar graphs showing the signal intensities of PG and BMP species in WT and CLN3-KO mice. Error bars represent the standard deviation of three biological replicates. Statistical analysis was performed using an unpaired two-tailed Student’s t-test (*P ≤ 0.05; ns, not significant). c) Mobility-constrained MALDI-iPRM ion images of PG 16:0_20:2, showing the precursor ion image at m/z 773.5337 and the corresponding product ion images at m/z 307.2637 and 255.2314. Mobility-constrained MALDI-iPRM images of BMP 18:1_18:1, showing the precursor ion image at m/z 773.5337 and the corresponding product ion image at m/z 281.2479. Scale bar, 1 mm. Pixel size, 20 μm; raster size, 40 μm. Data were normalized by RMS.

The detected lipids species characterized by a total of 36 carbon atoms and two double bonds provided a particularly informative example. Indeed, the as mobility selection of the common precursor at *m/z* 773.5337 distinguished two molecular species with different acyl-chain compositions. The PG-associated mobility region generated product ions at *m/z* 255.23 and 307.26, supporting its annotation as PG 16:0_20:2, whereas the BMP-associated fragments predominantly generated the 18:1 fatty-acyl anion at *m/z* 281.25, consistent with BMP 18:1_18:1 (**Figure 5c)**. Thus, mobility-constrained fragmentation not only discriminated the two isomeric lipid classes, but also revealed distinct molecular structures underlying the same sum composition. Complete MALDI-iPRM product-ion images are reported in **Figure S11**.

At the regional level, PG 34:1 displayed a structured renal distribution, with higher signal intensity in the cortex (Cx) and inner stripe of the outer medulla (ISOM) and lower signal toward the inner medulla compartment; this pattern was largely preserved in CLN3-KO tissue. In contrast, BMP 16:0_18:1 and BMP 18:1_18:1 showed a more widespread reduction in the KO kidney, without an evident restriction to a single anatomical compartment. Notably, these regional patterns were consistently reproduced in the corresponding fatty-acyl product-ion images, supporting the spatial concordance between mobility-selected precursor and product-ion distributions.

The applicability of the workflow was further demonstrated in lung, which yielded the largest number of detectable PG/BMP pairs among the tissues investigated. Mobility-resolved precursor and product-ion images supported the reciprocal detection of the 16:0_16:0, 16:0_16:1, 16:0_18:1, 16:0_18:2, 16:0_20:2, 18:1_18:1, 16:0_22:6 and 18:2_20:4 compositions (**Figure 6** and **Figure S12**). A similar genotype-dependent pattern was observed in lung tissue, where BMP species showed lower signal intensities in CLN3-KO mice, while PG signals remained comparable between genotypes (**Figure 6a,b; Figure S12**).

**Figure 6.**
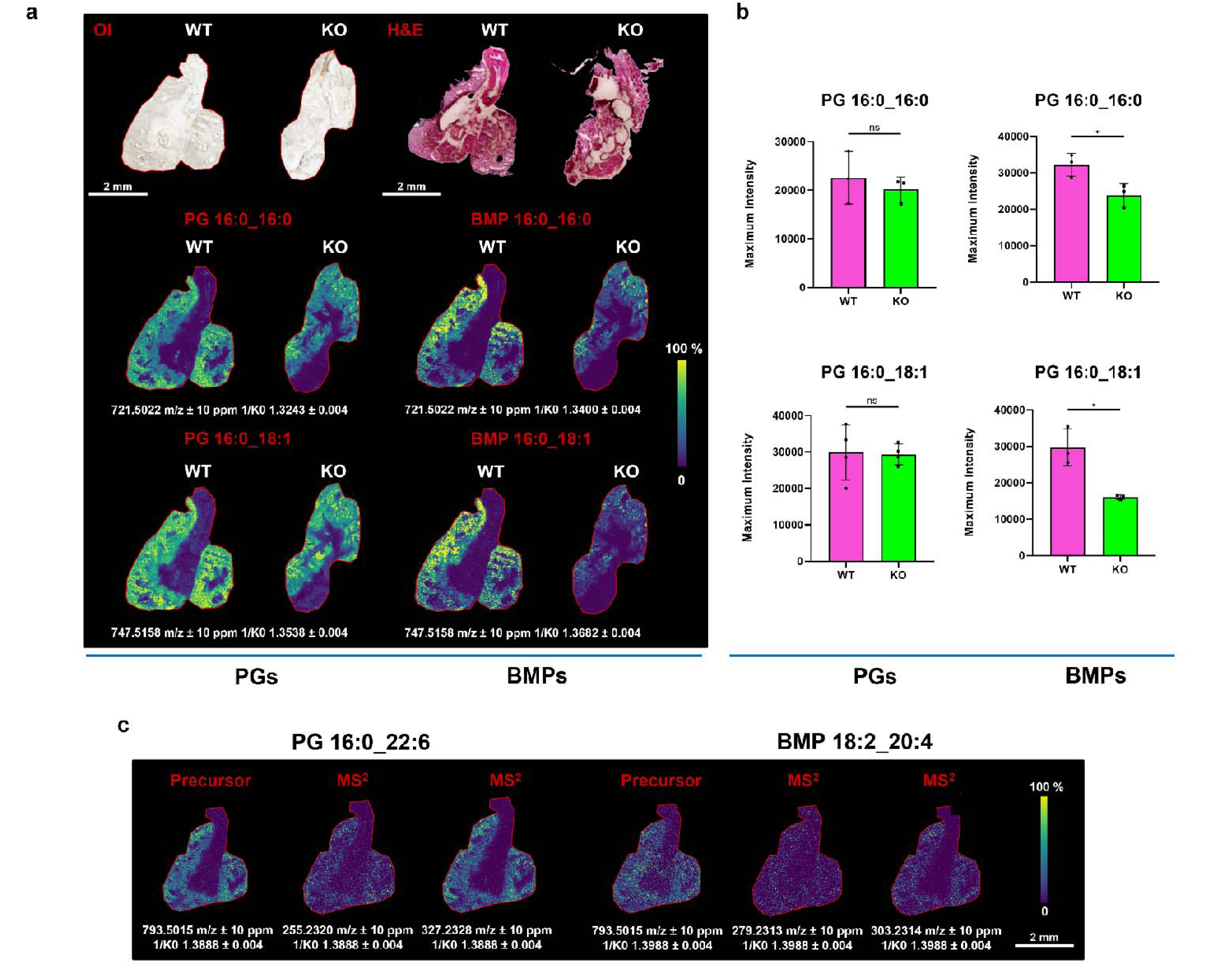
MALDI-iPRM discrimination of PGs and BMPs in lung sections from WT and CLN3-knockout (KO) mice. a) MALDI-TIMS MS^1^ ion images of PG/BMP 16:0_16:0 and PG/BMP 16:0_18:1 in lung sections from WT and CLN3-KO mice. Scale bar, 2 mm. Color scale bar is shown as percentage of maximum intensity. Abbreviations: OI, Optical Image; H&E, Hematoxylin and Eosin Staining. b) Bar graphs showing the signal intensities of PG and BMP species in WT and CLN3-KO mice. Error bars represent the standard deviation of three biological replicates. Statistical analysis was performed using an unpaired two-tailed Student’s t-test (*P ≤ 0.05; ns, not significant). c) Mobility-constrained MALDI-iPRM ion images of PG 16:0_22:6, showing the precursor ion image at m/z 793.5015 and the corresponding product ion images at m/z 255.2320 and 327.2328. Mobility-constrained MALDI-iPRM images of BMP 18:2_20:4, showing the precursor ion image at m/z 793.5015 and the corresponding product ion image at m/z 279.2313 and 303.2314. Scale bar, 1 mm. Pixel size, 20 μm; raster size, 40 μm. Data were normalized by RMS.

Importantly, several identical nominal PG/BMP elemental compositions corresponded to different acyl-chain combinations. For example, PG 38:6 was assigned as 16:0_22:6 from the product ions at *m/z* 255.23 and 327.23, whereas the corresponding BMP-associated population generated product ions at *m/z* 279.23 and 303.23, supporting BMP 18:2_20:4. These observations further illustrate the value of combining mobility-selective precursor isolation with spatially resolved MS/MS, since accurate mass or sum-composition annotation alone would not distinguish either the lipid class or the underlying fatty-acyl composition. Complete MALDI-iPRM product-ion images are reported in **Figure S13**.

The comparatively broad BMP coverage observed in lung is biologically noteworthy. BMPs are highly enriched in late endosomal and lysosomal membranes and have been reported at particularly high levels in lysosome-rich pulmonary cell populations, including alveolar macrophages ^40^. The diversity of BMP species detectable in lung is therefore consistent with its cellular and lysosomal composition. Nevertheless, the present MSI data do not provide cell-type-specific assignment and dedicated histological or molecular markers would be required to establish whether individual BMP distributions originate specifically from alveolar macrophage population.

## Conclusions

In this work, we developed an ion mobility-guided MALDI-iPRM-PASEF workflow for the reciprocal discrimination of endogenous BMP and PG isomers directly in tissue. Orthogonal HILIC-DDA-PASEF analysis provided the accurate-mass, fragmentation, and experimentally determined ion-mobility information required to define selective precursor coordinates for targeted MALDI-MS/MS imaging. Although PG and BMP species were not baseline-resolved by TIMS, optimization of ion load and mobility windows reduced reciprocal cross-talk to below 4% and enabled selective fragmentation of both isomeric populations. The workflow further provided acyl-chain-resolved precursor and product-ion imaging and resolved type-II isotopic interference that remained ambiguous at the MS¹ level. The analytical strategy proved transferable across brain, kidney, and lung tissues and enabled discrimination of endogenous PG/BMP pairs with different relative abundances and fatty-acyl compositions. Application to CLN3-knockout mice further illustrated its biological utility, revealing genotype-dependent reductions in BMP-associated signals that were not mirrored by the corresponding PG isomers. Importantly, the approach does not require complete baseline ion-mobility separation, provided that mobility-selective precursor isolation is combined with targeted product-ion imaging. The method nevertheless has some limitations. As a targeted workflow, it requires prior characterization of the relevant precursor *m/z*–mobility coordinates, here obtained by orthogonal HILIC-DDA-PASEF analysis. Ion-mobility measurements are also sensitive to acquisition conditions and ion load, requiring careful control of laser sampling to preserve predefined mobility coordinates. Structural characterization is currently achieved at the lipid-class and fatty-acyl composition level and does not provide *sn*-position or double-bond positional information. Absolute quantification was beyond the scope of the present study and would require dedicated isotope-labelled calibration, assessment of tissue-dependent matrix effects, and appropriate matrix-matched validation. Overall, the workflow provides a transferable analytical strategy for the confident spatial discrimination and structural confirmation of endogenous BMP/PG isomers in complex tissues.

## Supporting information

Supplementary Material_bioRxiv.docx

## Author Contributions

Conceptualization: E.So. and E.Sa.; implementation: E.So.; imaging experiments and data analysis: E.Sa., F.M; data interpretation: E.Sa., F.M., and D.L.M; funding acquisition: P.C.; drafting the manuscript: E.So., E.Sa. and D.L.M.; reviewing and editing the paper: D.L.M., P.C.

## Acknowledgments

This work was supported by Ministero dell’Università e della Ricerca (MIUR, Italy) project “Pathogen Readiness Platform for CERIC ERIC upgrade” - PRP@CERIC CUP J97G22000400006, project National Center for Gene Therapy and Drugs based on RNA Technology CUP: D43C2200120000, project PNC0000001 “D34 Health—Digital Driven Diagnostics, prognostics and therapeutics for sustainable Health care” to P. Campiglia. D.L.M is supported by the NCL Stiftung, Germany

**Supplementary Material_bioRxiv.docx**

