## Supplementary Material_bioRxiv.docx for "Ion Mobility-Guided Tandem Mass Spectrometry Imaging Resolves Bis(monoacylglycero)phosphate and Phosphatidylglycerol Isomers in Tissue"

**Table S1.** Application conditions for Norhamane (NRM) and naphthylethylenediamine dihydrochloride (NEDC) matrices.

| **MALDI Matrix** | **NRM** | **NEDC** |
| --- | --- | --- |
| **Concentration** | 7.5 mg/mL in 80% MeOH | 7 mg/mL in 80% MeOH |
| **Nozzle T (°C)** | 60 | 90 |
| **N_2_ Pressure (psi)** | 10 | 10 |
| **Flow rate (μL/min)** | 100 | 70 |
| **Nozzle velocity (mm/min)** | 1200 | 1200 |
| **Track spacing (mm)** | 2.5 | 2 |
| **Number of passes** | 15 | 6 |
| **Spray pattern** | Criss Cross (CC) | Criss Cross (CC) |

**Table S2**. Determination coefficient (R^2^), Limits of Detection (LODs) and Limits of Quantitation (LOQs) expressed as picomoles per mm² (pmol/mm²) of PG 18:1_18:1 and BMP 18:1_18:1 standards spotted onto kidney tissue sections. PG serial dilutions of PG and BMP standards were spotted at six concentration levels (0.1, 1.2, 2.9, 4.9, 11.5 and 20 pmol/mm²), together with a MeOH/H₂O (1:1, v/v) control. Calibration curves were generated by plotting the measured signal response against the amount of analyte deposited per unit area (pmol/mm²). Limits of detection (LODs) and quantification (LOQs) were calculated by the ratio between the standard deviation (σ) of the response at lowest analyte concentration and the slope of the calibration curve (S) multiplied by 3.3 and 10, respectively, according to the following formulae:

$$LOD= 3.3 \times\left( \frac{\sigma}{S} \right) LOQ= 10 \times(\frac{\sigma}{S})$$

|  | **Compound** |  | **R^2^** | **LODs (pmol/mm^2^)** | **LOQs (pmol/mm^2^)** |
| --- | --- | --- | --- | --- | --- |
|  | **PG 18:1_18:1** |  | **0.9937** | 0.430 | 1.433 |
|  | **BMP 18:1_18:1** |  | **0.9984** | 0.349 | 1.162 |

**Table S3.** List of PG, BMP, and LP species detected and annotated in lung, kidney, and brain tissues by HILIC-DDA-PASEF analysis

| **RT [min]** | **Mob. 1/K0** | **CCS (Å²)** | **m/z** | **Ions** | **Name** | **Molecular Formula** | **Δm/z [ppm]** | **ΔCCS [%]** | **Lung** | **Kidney** | **Brain** |
| --- | --- | --- | --- | --- | --- | --- | --- | --- | --- | --- | --- |
| 3.74 | 1.45 | 295.1 | 865.5027 | [M-H]- | BMP 22:6_22:6 | C50H75O10P | 0.197 | -0.1 | ✓ | ✓ | ✓ |
| 3.75 | 1.429 | 290.9 | 841.5025 | [M-H]- | BMP 20:4_22:6 | C48H75O10P | -0.046 | -0.1 | ✓ | ✓ | ✓ |
| 3.76 | 1.423 | 289.6 | 839.4866 | [M-H]- | BMP 20:5_22:6 | C48H73O10P | -0.311 | 0 | ✓ | ✓ | ✓ |
| 3.82 | 1.443 | 293.7 | 845.532 | [M-H]- | BMP 20:2_22:6 | C48H79O10P | -2.172 | 0.1 | ✓ | ✓ | ✓ |
| 3.83 | 1.472 | 299.4 | 873.5617 | [M-H]- | BMP 22:4/22:4 | C50H83O10P | -3.876 | -0.1 | ✓ | ✓ | ✓ |
| 3.87 | 1.413 | 287.8 | 817.5024 | [M-H]- | BMP 18:2_22:6 | C46H75O10P | -0.196 | -0.1 | ✓ | ✓ | ✓ |
| 3.87 | 1.512 | 307.5 | 905.6269 | [M-H]- | BMP 24:0_22:6 | C52H91O10P | -0.858 | -0.1 | ✓ | × | × |
| 3.89 | 1.422 | 289.6 | 819.5176 | [M-H]- | BMP 18:1_22:6 | C46H77O10P | -0.741 | -0.1 | ✓ | ✓ | ✓ |
| 3.92 | 1.433 | 291.8 | 823.5469 | [M-H]- | BMP 18:1_22:4 | C46H81O10P | -3.061 | -0.6 | ✓ | ✓ | ✓ |
| 3.92 | 1.395 | 284.3 | 793.5024 | [M-H]- | BMP 18:2_20:4 | C44H75O10P | -0.159 | -0.6 | ✓ | ✓ | ✓ |
| 3.93 | 1.403 | 285.8 | 795.5175 | [M-H]- | BMP 18:1_20:4 | C44H77O10P | -0.864 | 0.2 | ✓ | ✓ | ✓ |
| 4.2 | 1.403 | 285.9 | 807.5179 | [M-H]- | BMP 17:0_22:6 | C45H77O10P | -0.269 | -0.1 | ✓ | ✓ | ✓ |
| 4.2 | 1.417 | 288.6 | 799.5469 | [M-H]- | BMP 18:1_20:2 | C44H81O10P | -3.156 | -0.1 | ✓ | ✓ | ✓ |
| 4.2 | 1.412 | 287.7 | 797.5324 | [M-H]- | BMP 18:1_20:3 | C44H79O10P | -1.774 | -0.1 | ✓ | ✓ | ✓ |
| 4.2 | 1.368 | 279 | 767.4879 | [M-H]- | BMP 16:1_20:4 | C42H73O10P | 1.363 | -0.1 | ✓ | ✓ | ✓ |
| 4.21 | 1.378 | 280.9 | 783.5178 | [M-H]- | BMP 17:0_20:4 | C43H77O10P | -0.42 | -0.1 | ✓ | ✓ | ✓ |
| 4.21 | 1.369 | 279 | 781.5021 | [M-H]- | BMP 17:1_20:4 | C43H75O10P | -0.556 | -0.1 | ✓ | ✓ | ✓ |
| 4.23 | 1.376 | 280.5 | 769.5025 | [M-H]- | BMP 18:2_18:2 | C42H75O10P | 0.021 | 0.2 | ✓ | ✓ | ✓ |
| 4.25 | 1.412 | 287.7 | 811.5484 | [M-H]- | BMP 17:0_22:4 | C45H81O10P | -1.333 | -0.1 | × | ✓ | ✓ |
| 4.25 | 1.422 | 289.7 | 801.5631 | [M-H]- | BMP 18:1_20:1 | C44H83O10P | -2.524 | -0.1 | ✓ | ✓ | ✓ |
| 4.27 | 1.341 | 273.6 | 741.4716 | [M-H]- | BMP 16:1_18:3 | C40H71O10P | 0.548 | -0.1 | ✓ | × | ✓ |
| 4.27 | 1.393 | 284 | 773.5334 | [M-H]- | BMP 18:1_18:1 | C42H79O10P | -0.564 | -0.1 | ✓ | ✓ | ✓ |
| 4.27 | 1.478 | 300.8 | 857.6271 | [M-H]- | BMP 18:2_24:0 | C48H91O10P | -0.692 | -0.1 | ✓ | × | × |
| 4.29 | 1.349 | 275.2 | 743.4869 | [M-H]- | BMP 16:1_18:2 | C40H73O10P | 0.021 | -0.1 | ✓ | ✓ | ✓ |
| 4.3 | 1.358 | 277.1 | 745.502 | [M-H]- | BMP 16:1_18:1 | C40H75O10P | -0.723 | -0.1 | ✓ | ✓ | ✓ |
| 4.31 | 1.326 | 270.7 | 717.4708 | [M-H]- | BMP 16:1_16:1 | C38H71O10P | -0.645 | -0.1 | ✓ | × |  |
| 4.32 | 1.367 | 278.7 | 747.5164 | [M-H]- | BMP 16:0_18:1 | C40H77O10P | -2.371 | -0.2 | ✓ | ✓ | ✓ |
| 4.51 | 1.334 | 272.2 | 719.4864 | [M-H]- | BMP 16:0_16:1 | C38H73O10P | -0.635 | -0.7 | ✓ | ✓ | ✓ |
| 4.52 | 1.327 | 270.7 | 731.4867 | [M-H]- | BMP 15:0_18:2 | C39H73O10P | -0.254 | -0.1 | ✓ | ✓ | × |
| 4.52 | 1.344 | 274.3 | 721.5021 | [M-H]- | BMP 16:0_16:0 | C38H75O10P | -0.612 | 0.1 | ✓ | ✓ | ✓ |
| 4.53 | 1.302 | 265.8 | 705.4711 | [M-H]- | BMP 15:0_16:1 | C37H71O10P | -0.192 | -0.1 | ✓ | ✓ | × |
| 3.54 | 1.309 | 267.4 | 693.4716 | [M-H]- | BMP 14:0_16:0 | C36H71O10P | 0.609 | -0.2 | ✓ | × | × |
| 4.61 | 1.335 | 272.3 | 733.503 | [M-H]- | BMP 15:0_18:1 | C39H75O10P | 0.641 | -0.1 | ✓ | ✓ | ✓ |
| 4.61 | 1.394 | 283.9 | 805.5037 | [M-H]- | BMP 17:1_22:6 | C45H75O10P | 1.439 | -0.1 | × | ✓ | ✓ |
| 4.7 | 1.28 | 261.5 | 679.4557 | [M-H]- | BMP 13:0_16:0 | C35H69O10P | 0.261 | -0.2 | ✓ | × | × |
| 5.31 | 1.295 | 264.5 | 693.4709 | [M-H]- | PG 14:0_16:0 | C36H71O10P | -0.386 | -0.1 | ✓ | ✓ | ✓ |
| 5.32 | 1.285 | 262.5 | 691.4551 | [M-H]- | PG 14:0_16:1 | C36H69O10P | -0.651 | -0.2 | ✓ | × | × |
| 5.48 | 1.41 | 287.1 | 819.5174 | [M-H]- | PG18:1_22:6 | C46H77O10P | -0.98 | 0.6 | ✓ | ✓ | ✓ |
| 5.49 | 1.418 | 288.8 | 821.5334 | [M-H]- | PG 18:0_22:6 | C46H79O10P | -0.511 | 0.3 | ✓ | ✓ | ✓ |
| 5.49 | 1.401 | 285.4 | 817.5022 | [M-H]- | PG18:2_22:6 | C46H75O10P | -0.345 | 0.3 | ✓ | ✓ | ✓ |
| 5.52 | 1.382 | 281.7 | 795.5176 | [M-H]- | PG 16:0_22:5 | C44H77O10P | -0.651 | 0.1 | ✓ | ✓ | ✓ |
| 5.52 | 1.384 | 282 | 793.5027 | [M-H]- | PG 16:0_22:6 | C44H75O10P | 0.182 | 0.2 | ✓ | ✓ | ✓ |
| 5.52 | 1.399 | 285 | 797.5338 | [M-H]- | PG 18:0_20:4 | C44H79O10P | -0.063 | 0.2 | ✓ | ✓ | ✓ |
| 5.52 | 1.374 | 279.9 | 791.4876 | [M-H]- | PG16:1_22:6 | C44H73O10P | 0.883 | 0.3 | ✓ | ✓ | ✓ |
| 5.54 | 1.424 | 290 | 825.5633 | [M-H]- | PG 18:0_22:4 | C46H83O10P | -2.17 | 0.2 | ✓ | × | ✓ |
| 5.55 | 1.361 | 277.5 | 769.5027 | [M-H]- | PG 16:0_20:4 | C42H75O10P | 0.231 | 0.1 | ✓ | ✓ | ✓ |
| 5.56 | 1.35 | 275.2 | 767.4867 | [M-H]- | PG 16:1_20:4 | C42H73O10P | -0.214 | 0.1 | ✓ | × | ✓ |
| 5.61 | 1.379 | 281.1 | 783.5179 | [M-H]- | PG17:0_20:4 | C43H77O10P | -0.343 | 0.1 | ✓ | × | ✓ |
| 5.61 | 1.465 | 298.1 | 857.6271 | [M-H]- | PG 18:2_24:0 | C48H91O10P | -0.685 | -0.2 | ✓ | × | × |
| 5.68 | 1.404 | 286.1 | 799.5482 | [M-H]- | PG 18:0_20:3 | C44H81O10P | -1.639 | -0.1 | ✓ | ✓ | ✓ |
| 5.69 | 1.472 | 299.5 | 859.6421 | [M-H]- | PG 18:1_24:0 | C48H93O10P | -1.516 | 0.1 | ✓ | × | × |
| 5.7 | 1.443 | 293.8 | 831.6109 | [M-H]- | PG22:0_18:1 | C46H89O10P | -1.393 | -0.1 | ✓ | × | × |
| 5.71 | 1.409 | 287.1 | 801.564 | [M-H]- | PG 18:0_20:2 | C44H83O10P | -1.365 | -0.1 | ✓ | ✓ | ✓ |
| 5.71 | 1.372 | 279.6 | 771.5178 | [M-H]- | PG 18:1_18:2 | C42H77O10P | -0.488 | -0.1 | ✓ | ✓ | ✓ |
| 5.71 | 1.38 | 281.2 | 773.5337 | [M-H]- | PG16:0_20:2 | C42H79O10P | -0.129 | -0.2 | ✓ | ✓ | ✓ |
| 5.81 | 1.329 | 271 | 743.4867 | [M-H]- | PG 16:1_18:2 | C40H73O10P | -0.248 | 0.1 | ✓ | ✓ | ✓ |
| 5.82 | 1.336 | 272.6 | 745.5025 | [M-H]- | PG 16:0_18:2 | C40H75O10P | -0.058 | -0.1 | ✓ | ✓ | ✓ |
| 5.82 | 1.374 | 280.2 | 761.5333 | [M-H]- | PG17:0_18:1 | C41H79O10P | -0.733 | -0.1 | ✓ | ✓ | ✓ |
| 5.83 | 1.354 | 276.1 | 747.5184 | [M-H]- | PG 16:0_18:1 | C40H77O10P | 0.28 | -0.1 | ✓ | ✓ | ✓ |
| 5.85 | 1.316 | 268.6 | 719.4864 | [M-H]- | PG 16:0_16:1 | C38H73O10P | -0.664 | 0.1 | ✓ | ✓ | ✓ |
| 5.86 | 1.349 | 275.2 | 735.5166 | [M-H]- | PG16:0_17:0 | C39H77O10P | -2.182 | 0.8 | ✓ | ✓ | ✓ |
| 5.9 | 1.326 | 270.6 | 721.5025 | [M-H]- | PG16:0_16:0 | C38H75O10P | 0.027 | 0.1 | ✓ | ✓ | ✓ |
| 6.11 | 1.307 | 266.9 | 707.4866 | [M-H]- | PG15:0_16:0 | C37H73O10P | -0.327 | -0.2 | ✓ | ✓ | ✓ |
| 6.11 | 1.4 | 285.3 | 807.5189 | [M-H]- | PG17:0_22:6 | C45H77O10P | 0.852 | 0.1 | ✓ | × | × |
| 7.12 | 1.055 | 217.3 | 483.2728 | [M-H]- | LPG 16:0 | C22H45O9P | -0.133 | -0.5 | ✓ | × | ✓ |
| 7.12 | 1.128 | 231.5 | 559.3037 | [M-H]- | LPG 22:4 | C28H49O9P | -0.738 | -0.1 | ✓ | ✓ | ✓ |
| 7.13 | 1.091 | 224.2 | 531.2729 | [M-H]- | LPG 20:4 | C26H45O9P | 0.189 | -0.1 | ✓ | ✓ | ✓ |
| 7.14 | 1.118 | 229.7 | 537.3191 | [M-H]- | LPG 20:1 | C26H51O9P | -1.334 | 0.1 | ✓ | ✓ | ✓ |
| 7.21 | 1.104 | 226.7 | 535.3021 | [M-H]- | LPG 20:2 | C26H49O9P | -3.773 | 0.1 | ✓ | ✓ | ✓ |
| 7.22 | 1.079 | 221.8 | 509.2883 | [M-H]- | LPG 18:1 | C24H47O9P | -0.416 | 0.1 | ✓ | ✓ | ✓ |
| 7.23 | 1.065 | 219.1 | 507.2728 | [M-H]- | LPG 18:2 | C24H45O9P | -0.192 | -0.1 | ✓ | ✓ | ✓ |
| 7.31 | 1.041 | 214.4 | 481.2568 | [M-H]- | LPG 16:1 | C22H43O9P | -0.777 | -0.1 | ✓ | ✓ | ✓ |
| 7.49 | 1.091 | 224.2 | 531.2725 | [M-H]- | LPG 20:4 | C26H45O9P | -0.695 | -0.1 | ✓ | ✓ | ✓ |
| 7.49 | 1.094 | 224.9 | 511.3037 | [M-H]- | LPG 18:0 | C24H49O9P | -0.894 | -0.3 | ✓ | ✓ | ✓ |
| 7.5 | 1.08 | 222 | 509.2882 | [M-H]- | LPG 18:1 | C24H47O9P | -0.603 | 0.1 | ✓ | ✓ | ✓ |
| 7.51 | 1.067 | 219.6 | 507.2726 | [M-H]- | LPG 18:2 | C24H45O9P | -0.549 | 0.1 | ✓ | ✓ | ✓ |
| 7.52 | 1.042 | 214.6 | 481.257 | [M-H]- | LPG 16:1 | C22H43O9P | -0.471 | -0.1 | ✓ | ✓ | ✓ |

**Table S4.** Comparison between theoretical and measured PG/BMP 18:1_18:1 ratios obtained after mobility-based signal extraction.

| **Standard Mixture** | **Theorical Ratio** | **Measured Ratio** |
| --- | --- | --- |
| **PG:BMP** | 1:1 | 1.12:1 ± 0.11 |
| **PG:BMP** | 10:1 | 10.44:1 ± 0.12 |
| **PG:BMP** | 50:1 | 31.57:1 ± 0.93 |
| **BMP:PG** | 10:1 | 8.92:1 ± 0.43 |

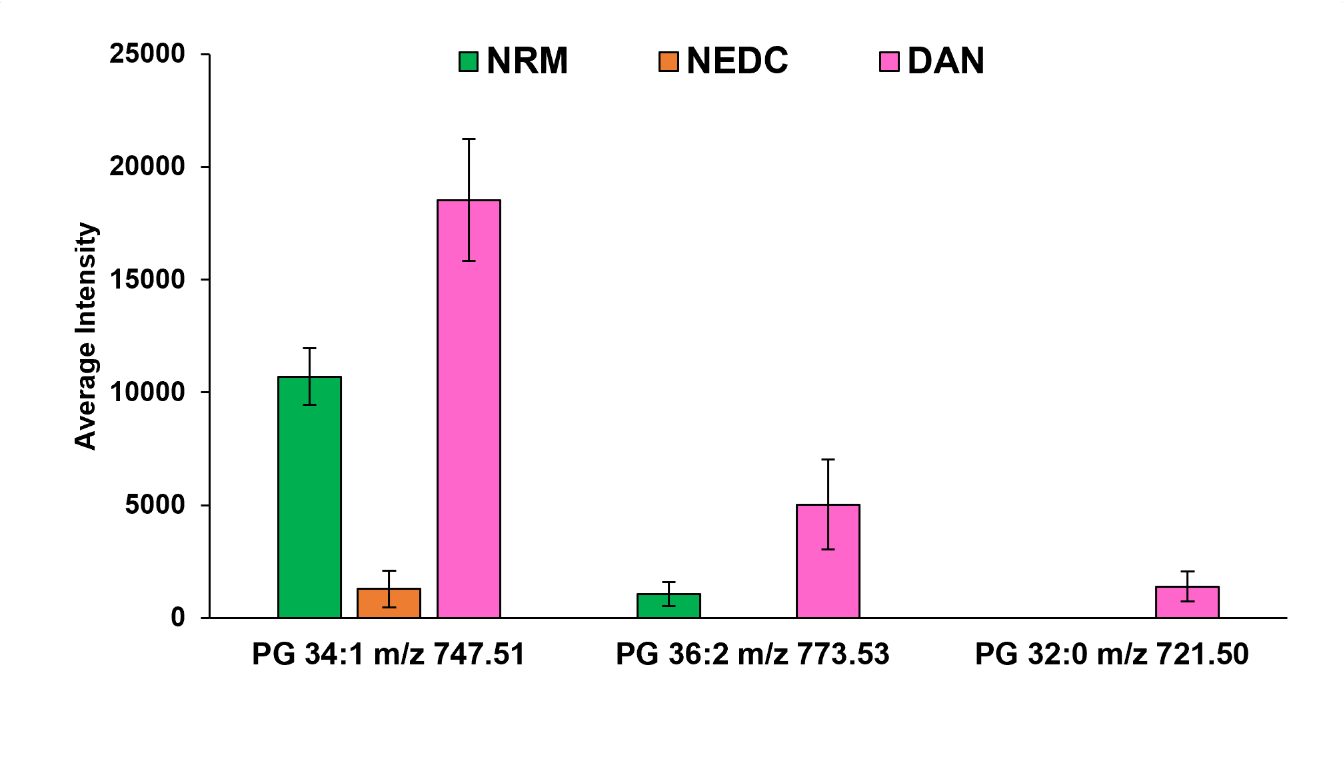

**Figure S1.** **Effect of MALDI matrix selection on PG detection in mouse kidney sections.** Bar graphs show the normalized average signal intensities of representative putatively annotated PG species detected in mouse kidney sections analyzed using different MALDI matrices, NRM, NEDC, and DAN, in negative-ion mode MALDI-TIMS-MSI.

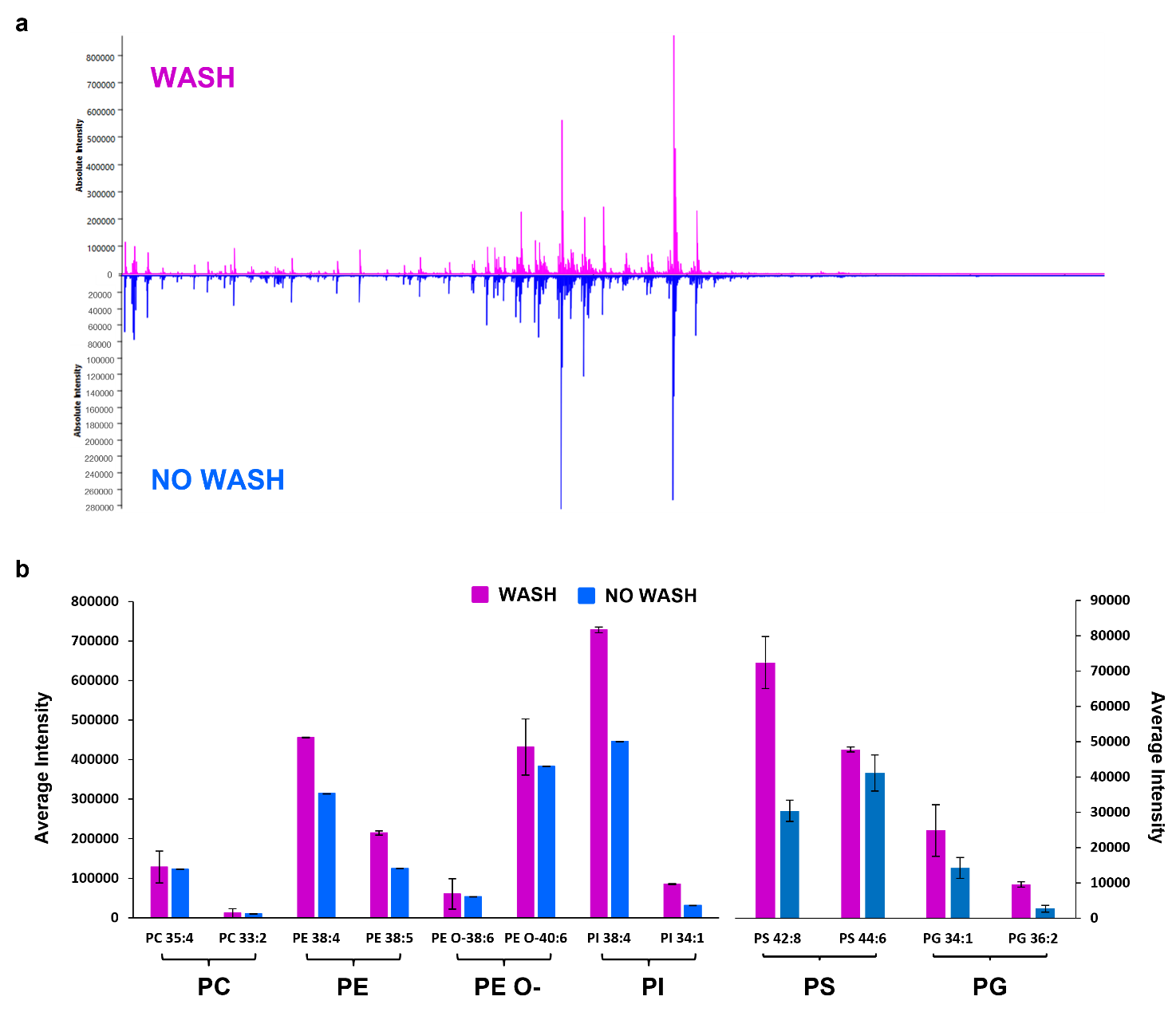

**Figure S2. Effect of ammonium acetate washing on lipid ionization in mouse kidney sections.** a) Representative Mean Spectrum profiles obtained from mouse kidney sections with and without a 50 mM CH_3_COONH_4_ washing step prior to matrix application. b) Bar graphs showing the normalized average signal intensities of representative lipid species from different lipid classes under washing and no-washing conditions.

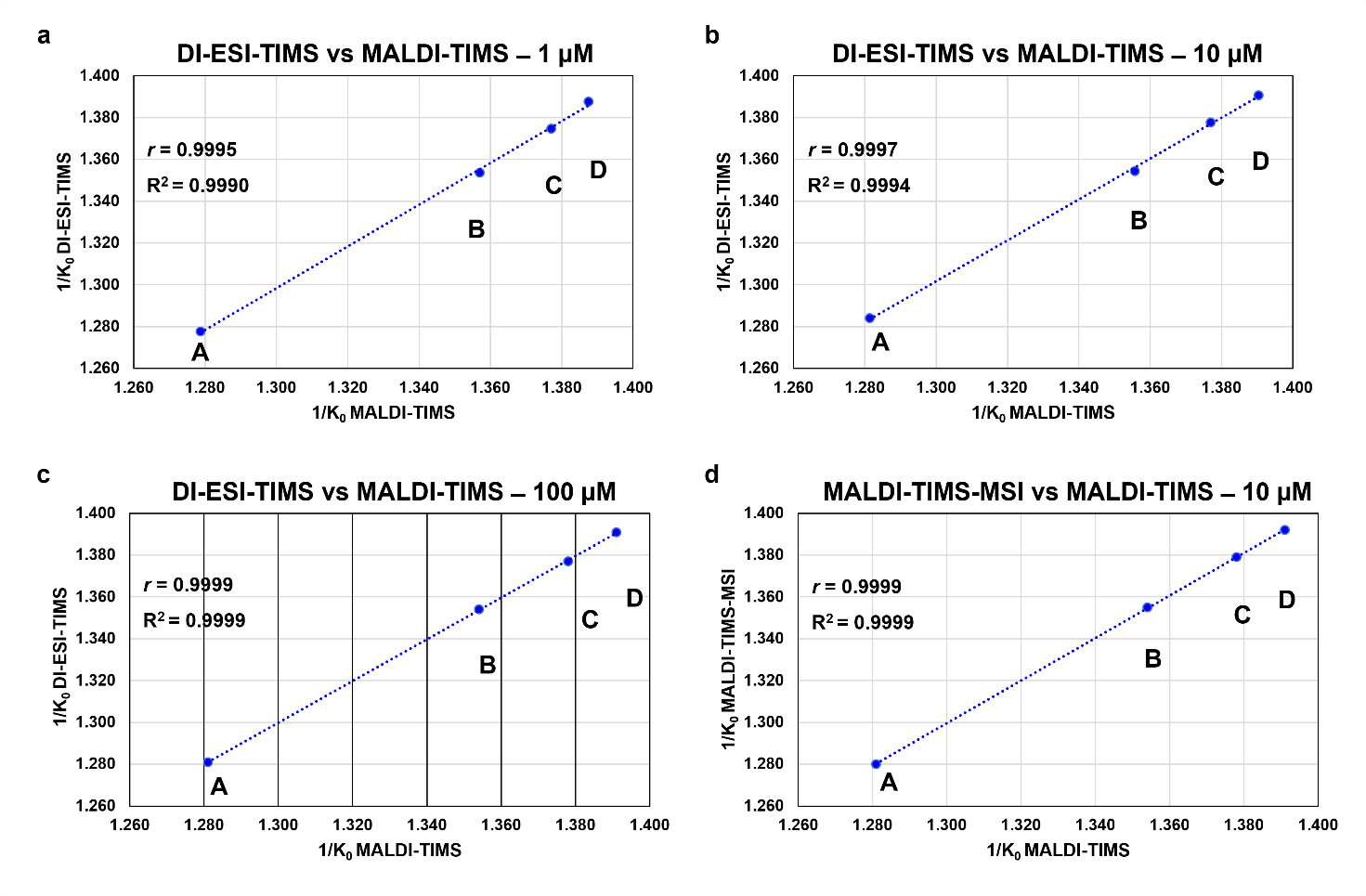

**Figure S3.** **Cross-platform correlation of ion mobility measurements.** a-c) Correlation plots comparing 1/K_0_ values measured by DI-ESI-TIMS and MALDI-TIMS for BMP 14:0_14:0 (A), PG 16:0_18:1 (B), PG 18:1_18:1 (C), BMP 18:1_18:1 (D) standards analyzed at 1, 10, and 100 µM. d) Correlation between 1/ K_0_ values obtained by MALDI-TIMS and tissue-based MALDI-TIMS-MSI for PG and BMP standards at concentration 10 µM. Pearson's correlation coefficient (*r*) and coefficient of determination (R^2^) are reported for each concentration level.

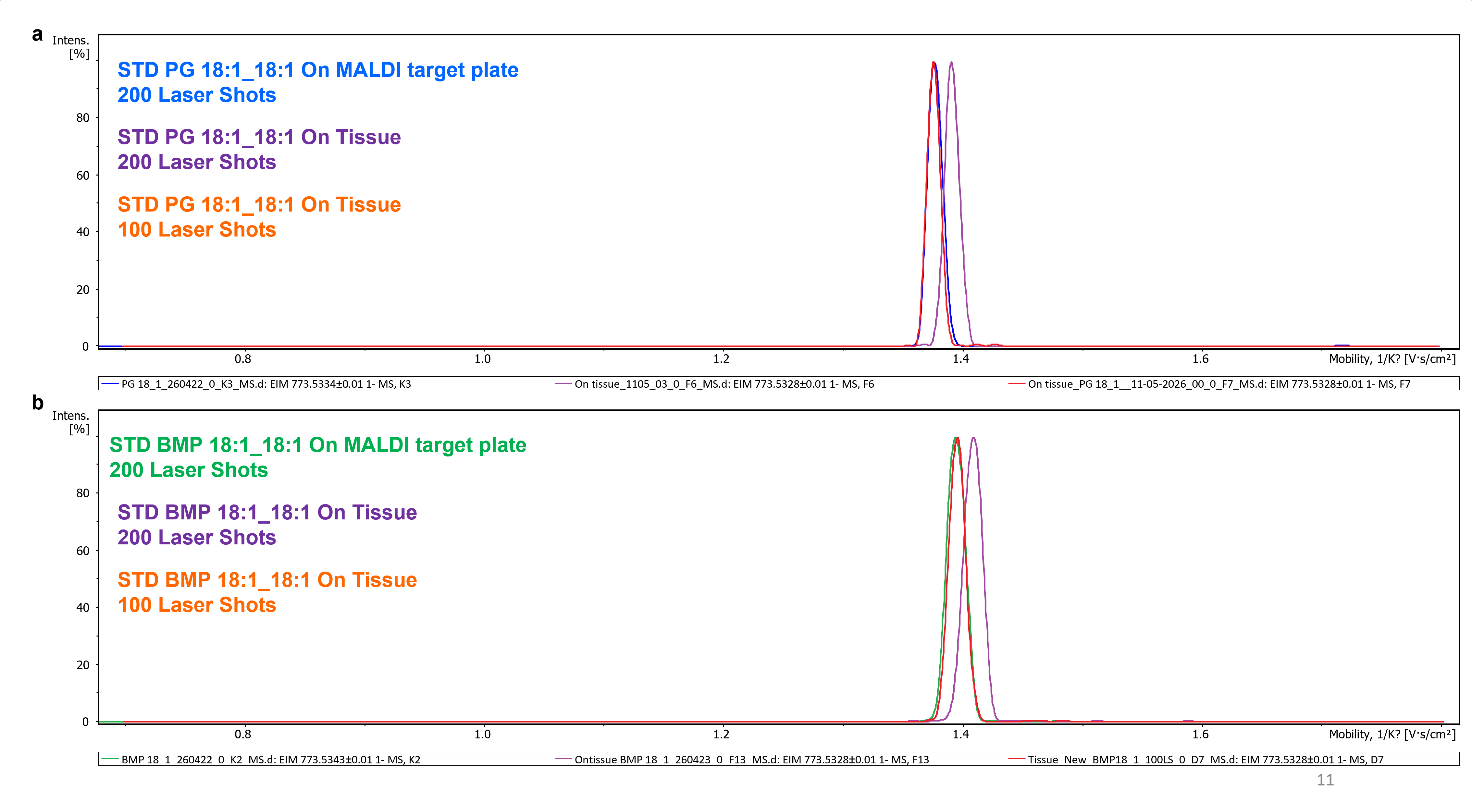

**Figure S4.** **Effect of laser shots on the ion mobility values 1/K_0_ of PG and BMPs species**. Extracted ion mobilograms (EIMs) of PG 18:1_18:1 (a) and BMP 18:1_18:1 (b) obtained from standards spotted on the MALDI target plate and on tissue sections. Measurements were performed using 100 and 200 laser shots.

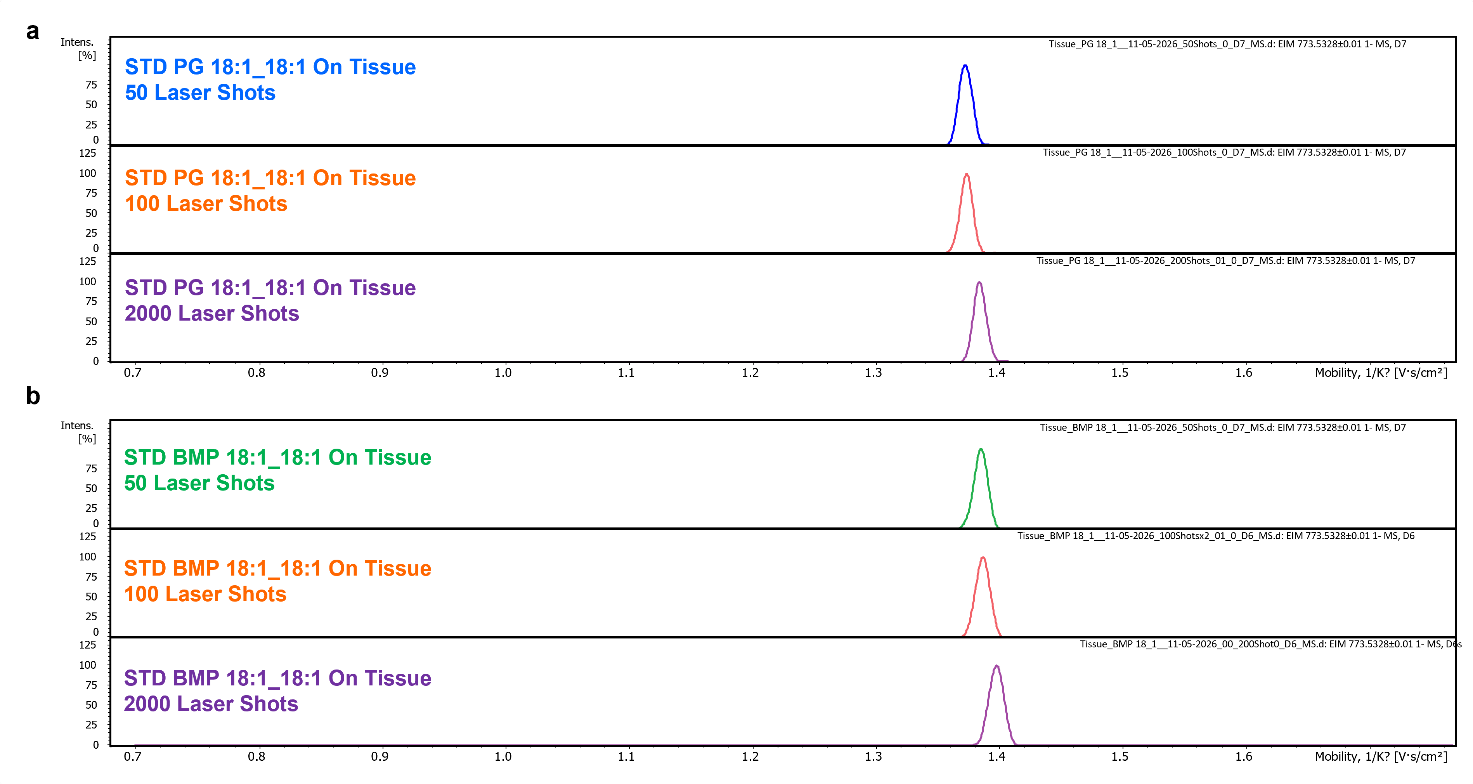

**Figure S5.** Extracted ion mobilograms EIMs showing the influence of increasing laser shots (50, 100, and 200) on the measured ion mobolity 1/K_0_ of a) PG 18:1_18:1 and b) BMP 18:1_18:1 standards spotted on tissue sections.

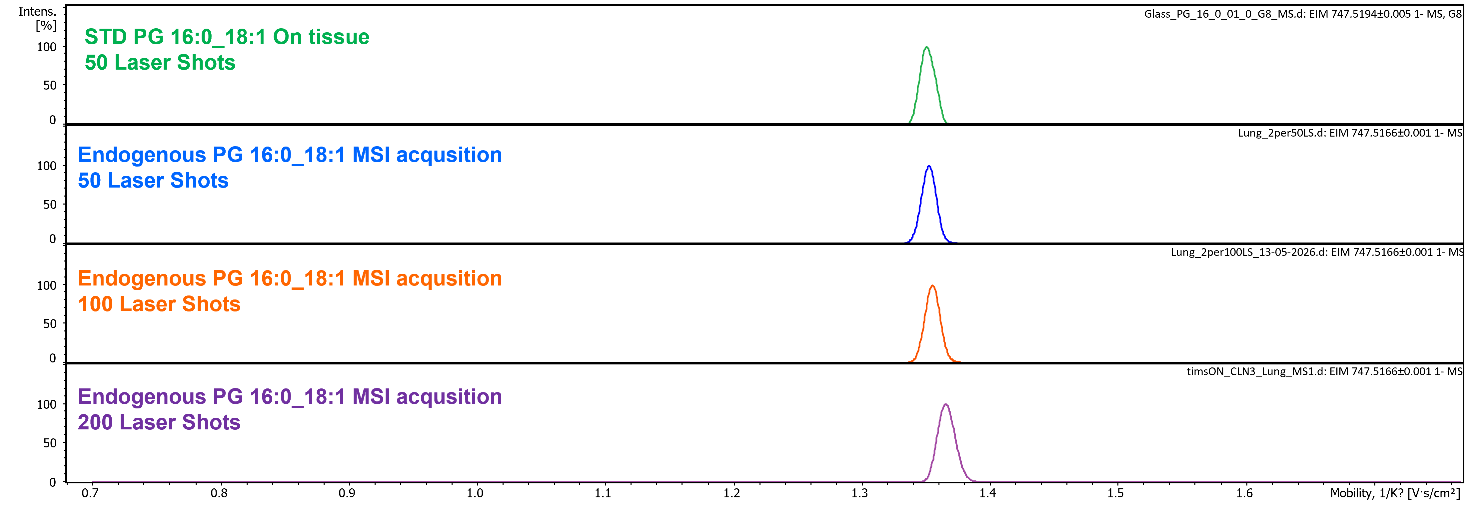

**Figure S6.** **Comparison of ion mobility distributions of endogenous and standard PG 16:0_18:1 under different laser shot conditions during MALDI-TIMS-MSI acquisition.** Extracted ion mobilograms (EIMs) of PG 16:0_18:1 obtained from endogenous tissue signal and from the corresponding pure standard spotted onto tissue sections using increasing numbers of laser shots (50, 100, and 200).

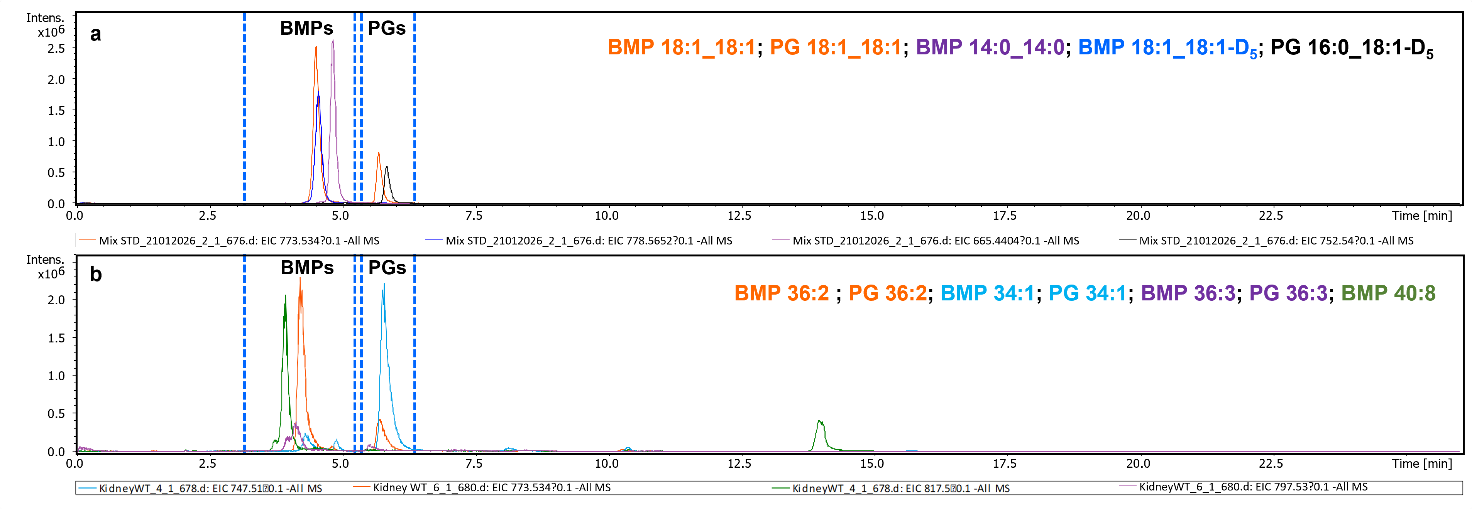

**Figure S7.** **HILIC-based separation and annotation of PG and BMP species.** a) Extracted ion chromatograms (EICs) of a mixture of authentic PG and BMP standards and deuterated internal standards (PG 18:1_18:1, BMP 18:1_18:1, BMP 14:0_14:0 (S,R), BMP 18:1_18:1-D_5_, PG 16:0-18:1-D_5_). b) EICs of endogenous PG and BMP species detected in a mouse kidney tissue extract. Highlighted retention time windows indicate the chromatographic regions corresponding to BMP and PG species and their co-elution with the deuterated internal standards, supporting the annotation of the endogenous lipid species.

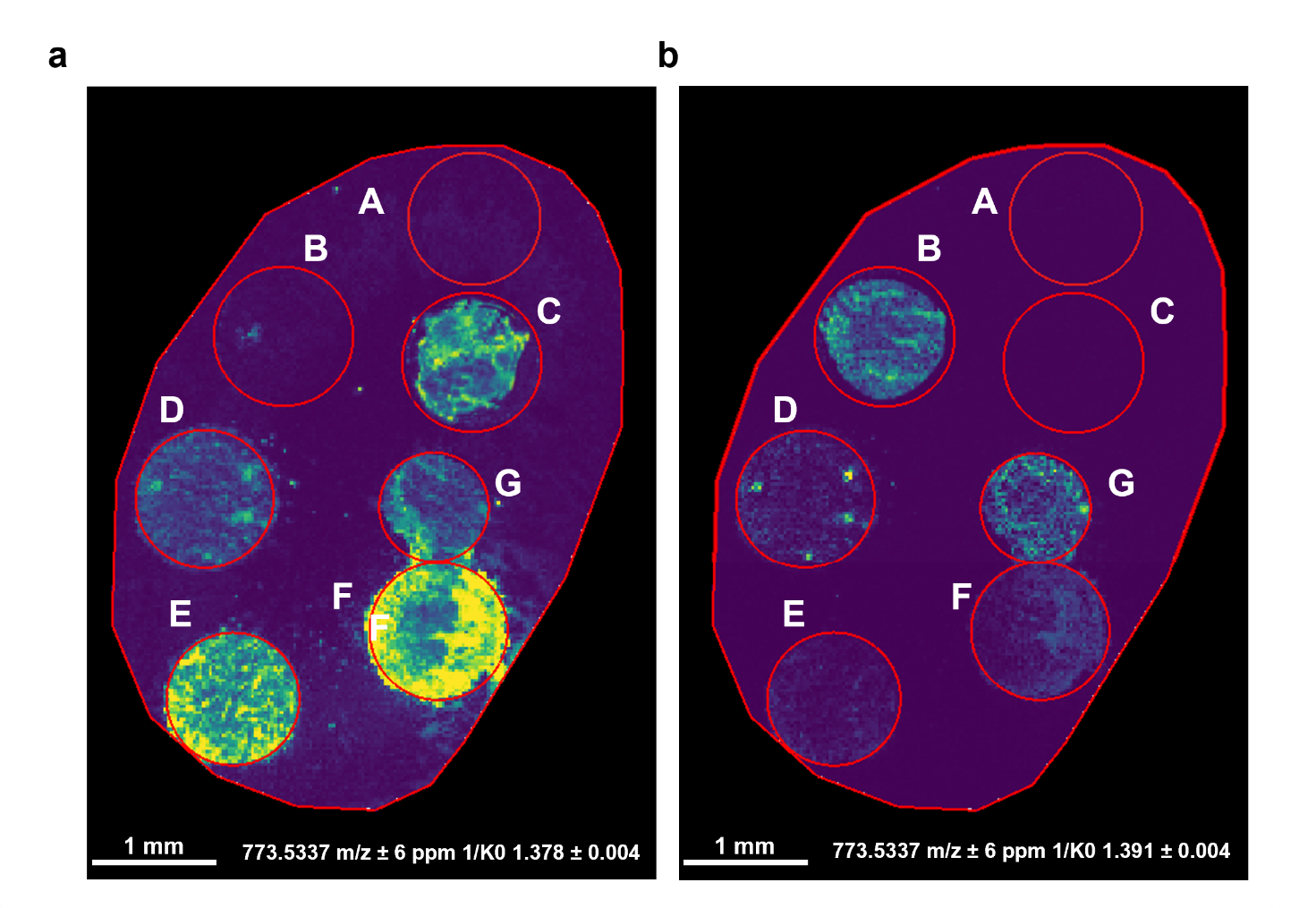

**Figure S8.** **MALDI-TIMS-MSI analysis of PG/BMP 18:1_18:1 standard mixtures spotted onto tissue sections.** Representative ion images obtained after mobility-based signal extraction using the characteristic 1/K_0_ values of PG and BMP (±0.004 1/K_0_). PG and BMP distributions within the mixed standard spots were independently visualized using their respective mobility extraction windows. Panel **a** shows the PG 18:1_18:1 distribution, panel **b** shows the BMP 18:1_18:1 distribution. Spot labels correspond to: (A) Blank MeOH/H₂O (1:1, v/v), (B) BMP 18:1_18:1 single standard (15 µM; 1.3 pmol/mm²), (C) PG 18:1_18:1 single standard (15 µM; 1.3 pmol/mm²), (D) PG/BMP 1:1, (E) PG/BMP 10:1, (F) PG/BMP 50:1, and (G) BMP/PG 10:1.

Scale bar = 1 mm. Color scale is expressed as percentage of the maximum signal intensity. Pixel size, 20 μm.

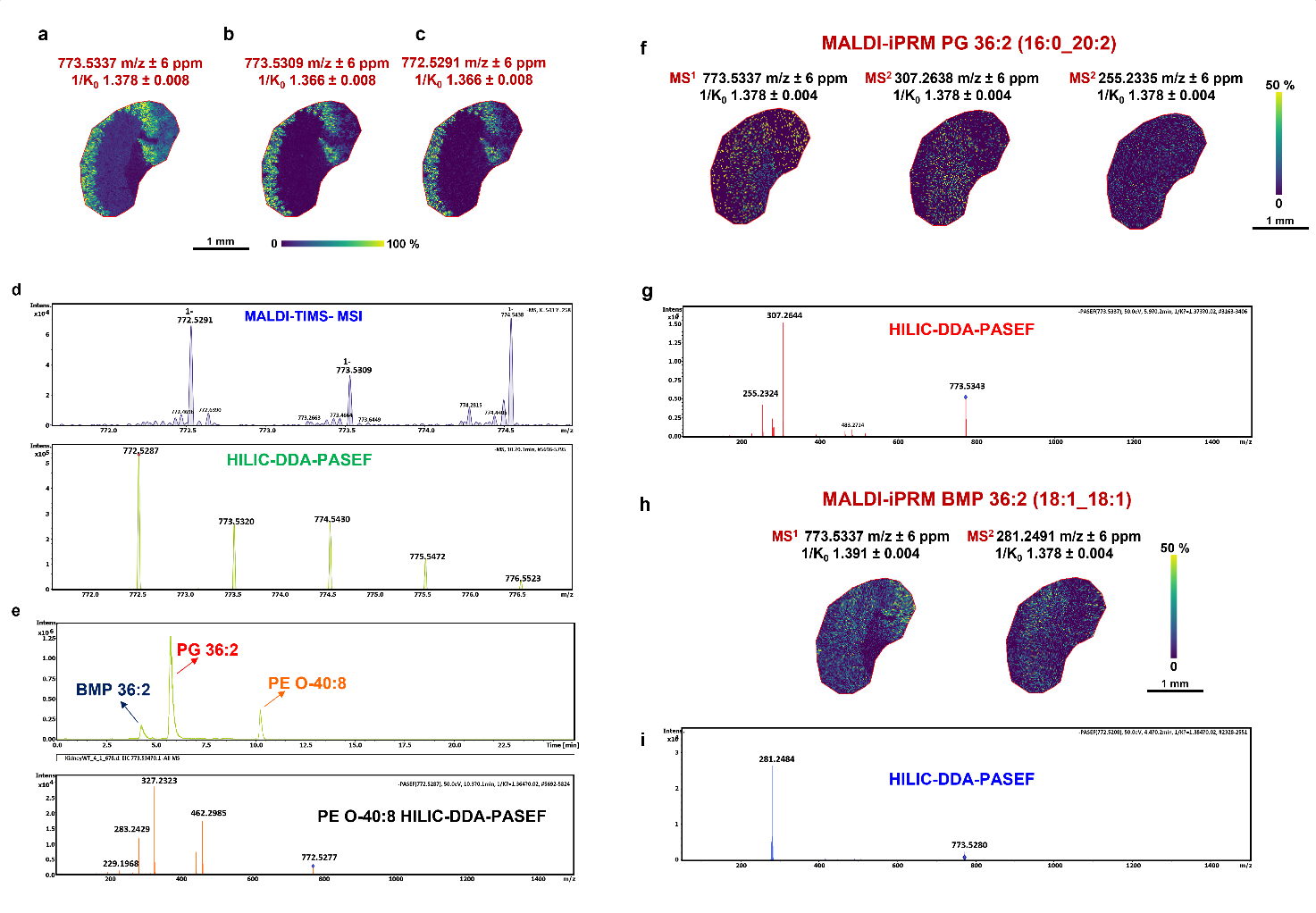

**Figure S9. Application of MALDI-iPRM for resolving isotopic interference affecting PG 36:2 annotation in kidney tissue sections.**

a-c) MALDI-TIMS-MS ion images showing the distribution of the precursor signal at m/z 773.5337 assigned to PG 36:2 (a), the spatially overlapping contribution arising from the M+1 isotopologue at m/z 773.5309 (b), and the corresponding precursor ion at m/z 772.5291 assigned to the co-distributed PE O-40:8 species (c).

d) Isotopic pattern demonstrating the partial contribution of the PE O-40:8 M+1 isotopologue within the extraction window selected for PG 36:2 for both MALDI-MSI (top) and HILIC-DDA-PASEF (bottom) spectra.

e) Extracted ion chromatogram (EIC) at m/z 773.5 showing the retention time of BMP/PG 36:2 and the M+1 isotopologue of PE O-40:8 (top), and the MS/MS spectrum of PE O-40:8 (bottom), obtained by HILIC-DDA-PASEF analysis.

f) MALDI-iPRM spectrum of the mobility-selected PG 36:2 precursor showing characteristic PG fragment ions, supporting its structural annotation as PG 16:0_20:2 and consistent with the identification obtained by HILIC-DDA-PASEF analysis (g).

h) MALDI-iPRM characterization of BMP 36:2 showing selective precursor isolation based on ion mobility and structural annotation as BMP 18:1_18:1, supported by HILIC-DDA-PASEF data (i).

Scale bar, 2 mm. Color scale bar is shown as percentage of maximum intensity. Pixel size, 20 μm; raster size, 40 μm. Data were normalized by RMS.

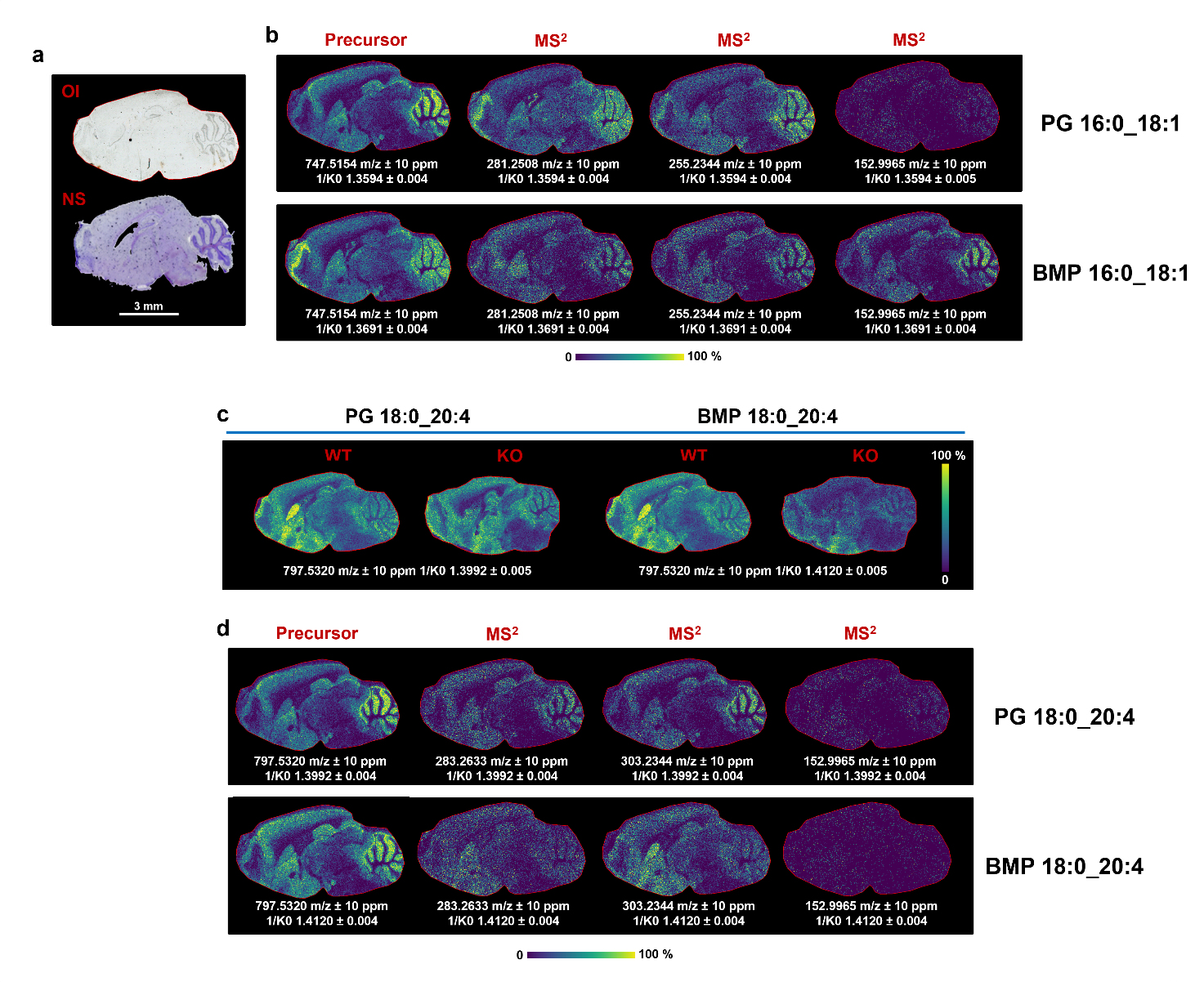

**Figure S10. MALDI-iPRM discrimination of PGs and BMPs in sagittal brain sections from WT and CLN3-knockout (KO) mice**.

a) Optical image (OI) and corresponding Nissl-stained section (NS).

b) MALDI-iPRM product ion images of PG 16:0_18:1 (top) and BMP 16:0_18:1 (bottom), generated from the mobility-selected precursor at m/z 747.5154 and the corresponding MS/MS fragment ions (MS^2^) at m/z 281.2508, 255.2344, and 152.9965.

c) MALDI-TIMS MS^1^ ion images of PG/BMP 18:0_20:4 in brain sections from WT and CLN3-KO mice.

d) MALDI-iPRM product ion images of PG 18:0_20:4 (top) and BMP 18:0_20:4 (bottom), generated from the mobility-selected precursor at m/z 797.5320 and the corresponding MS/MS fragment ions (MS^2^) at m/z 283.2633, 303.2344, and 152.9965.

Scale bar, 3 mm. Color scale bar is shown as percentage of maximum intensity. Pixel size, 20 μm; raster size, 40 μm. Data were normalized by RMS.

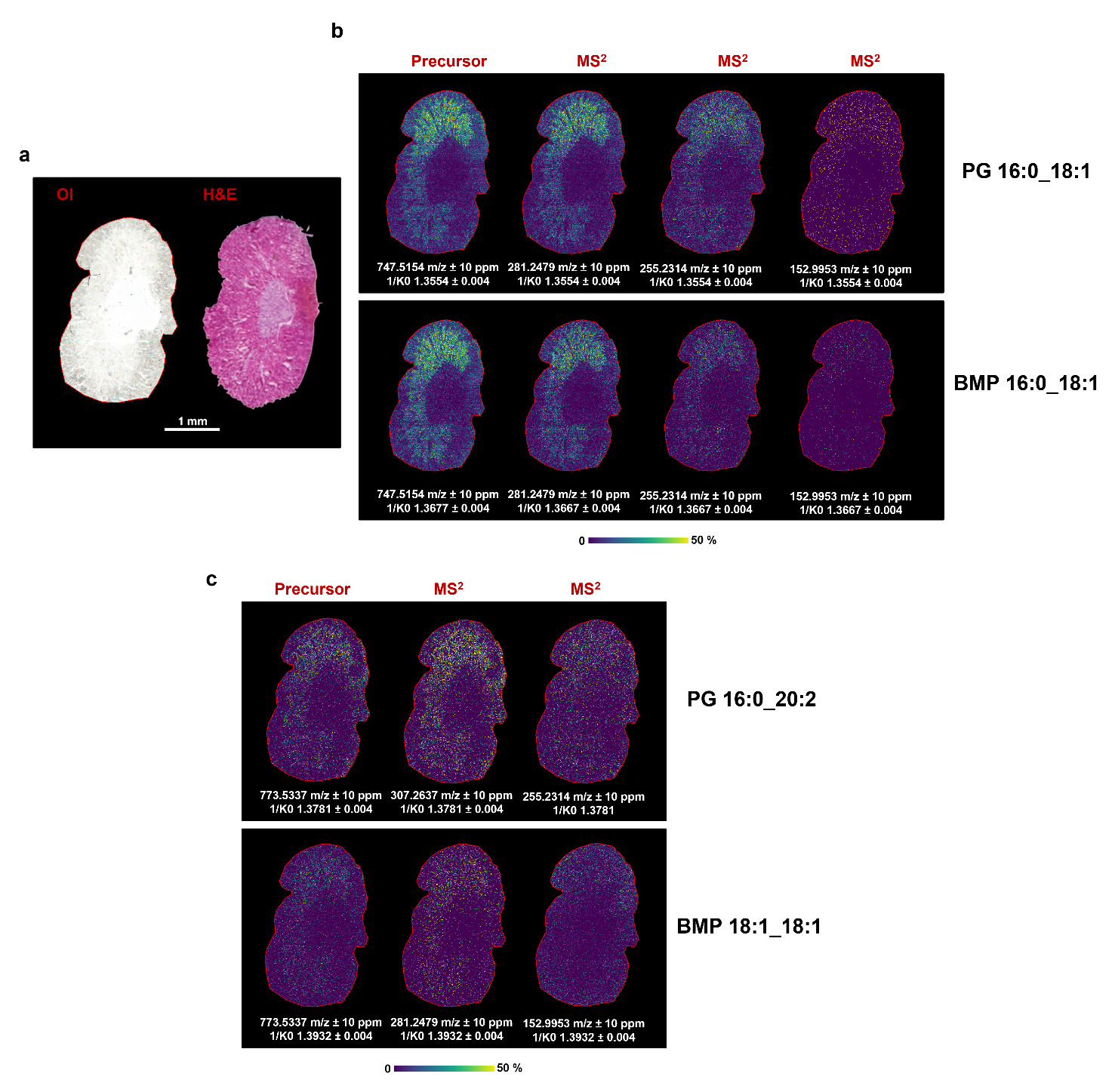

**Figure S11. MALDI-iPRM discrimination of PGs and BMPs in mouse kidney sections**.

a) Optical image (OI) and corresponding Hematoxylin and Eosin-stained section (H&E).

b) MALDI-iPRM product ion images of PG 16:0_18:1 (top) and BMP 16:0_18:1 (bottom), generated from the mobility-selected precursor at m/z 747.5154 and the corresponding MS/MS fragment ions (MS^2^) at m/z 281.2479, 255.2314, and 152.9953.

c) Mobility-constrained MALDI-iPRM ion images of PG 16:0_20:2 (top), showing the precursor ion image at m/z 773.5337 and the corresponding product ion images at m/z 307.2637 and 255.231, and of BMP 18:1_18:1 (bottom), showing the precursor ion image at m/z 773.5337 and the corresponding product ion image at m/z 281.2479 and 152.9953.

Scale bar, 1 mm. Color scale bar is shown as percentage of maximum intensity. Pixel size, 20 μm; raster size, 40 μm. Data were normalized by RMS.

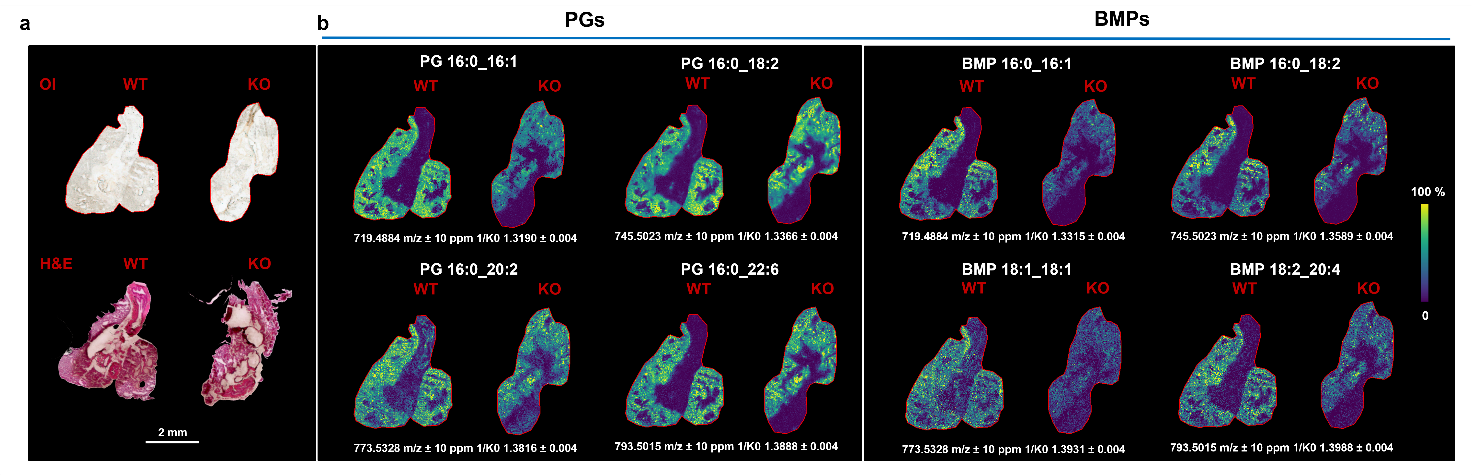

**Figure S12.** **MALDI-TIMS-MSI discrimination of PGs and BMPs in lung sections from WT and CLN3-knockout (KO) mice.**

a) Optical image (OI) and corresponding Hematoxylin and Eosin-stained sections (H&E).

b) MALDI-TIMS MS^1^ ion images of PG/BMP 16:0_16:1, PG/BMP 16.0_18:2, PG 16:0_20:2, BMP 18:1_18:1, PG 16:0_22:6 and BMP 18:2_20:4 in lung sections from WT and CLN3-KO mice. Scale bar, 2 mm. Color scale bar is shown as percentage of maximum intensity. Pixel size, 20 μm; raster size, 40 μm. Data were normalized by RMS.

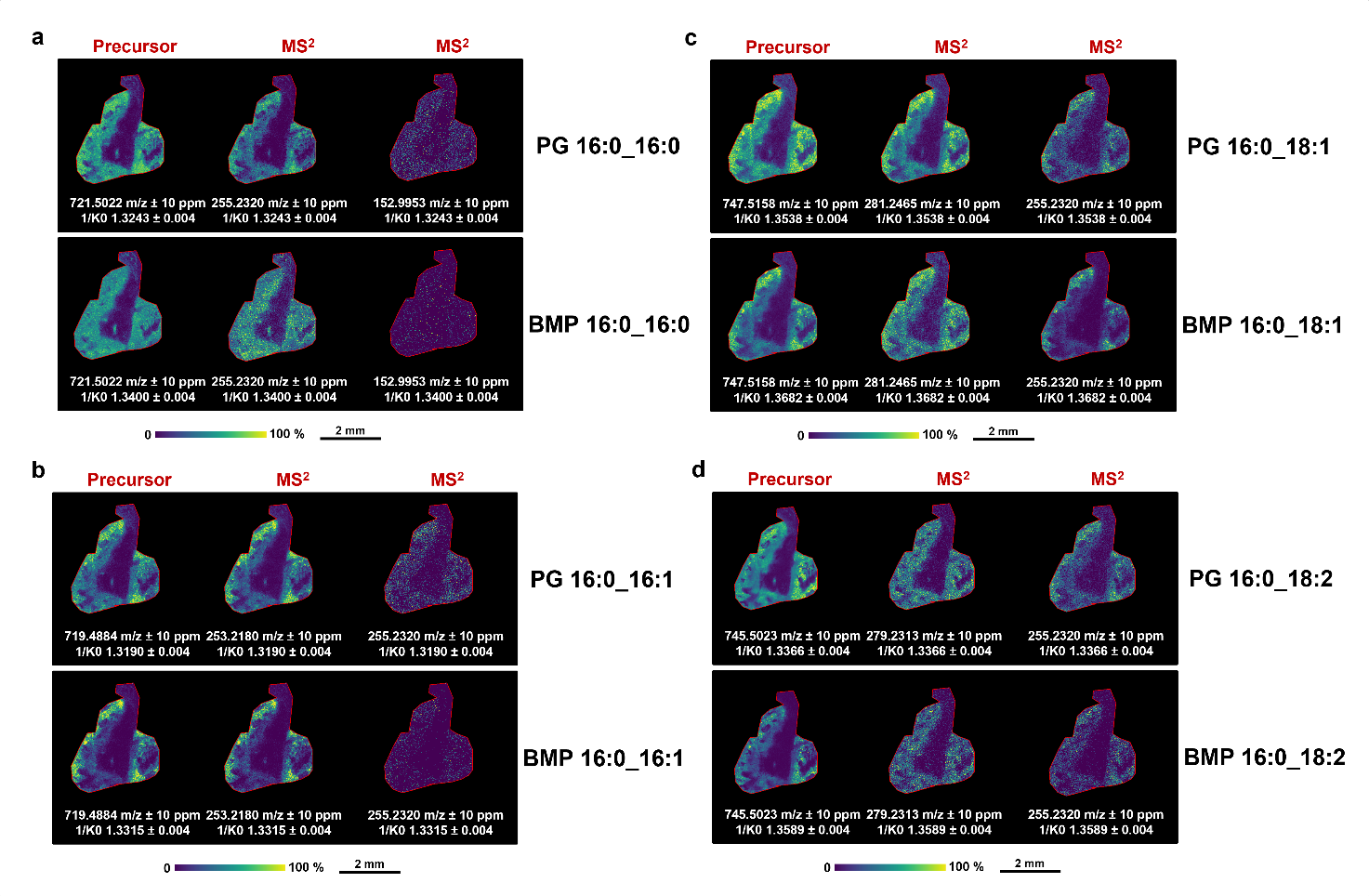

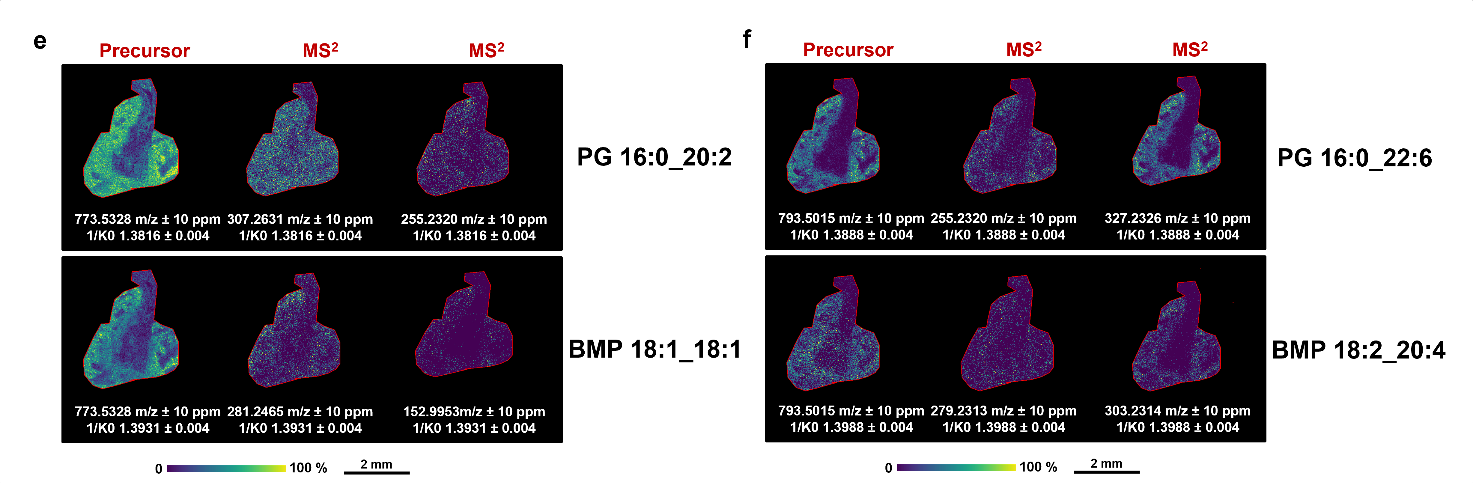

**Figure S13. MALDI-iPRM discrimination of PGs and BMPs in mouse lung sections.**

a) MALDI-iPRM product ion images of PG 16:0_16:0 (top) and BMP 16:0_16:0 (bottom), generated from the mobility-selected precursor at m/z 721.5022 and the corresponding MS/MS fragment ions (MS^2^) at m/z 255.2320 and 152.9953.

b) MALDI-iPRM product ion images of PG 16:0_16:1 (top) and BMP 16:0_16:1 (bottom), generated from the mobility-selected precursor at m/z 719.4884 and the corresponding MS/MS fragment ions (MS^2^) at m/z 253.2180 and 255.2320.

c) MALDI-iPRM product ion images of PG 16:0_18:1 (top) and BMP 16:0_18:1 (bottom), generated from the mobility-selected precursor at m/z 747.5158 and the corresponding MS/MS fragment ions (MS^2^) at m/z 281.2465 and 255.2320. d) MALDI-iPRM product ion images of PG 16:0_18:2 (top) and BMP 16:0_18:2 (bottom), generated from the mobility-selected precursor at m/z 745.5023 and the corresponding MS/MS fragment ions (MS^2^) at m/z 279.2313 and 255.2320.

d) Mobility-constrained MALDI-iPRM ion images of PG 16:0_20:2 (top), showing the precursor ion image at m/z 773.5337 and the corresponding product ion images at m/z 307.2637 and 255.231, and of BMP 18:1_18:1 (bottom), showing the precursor ion image at m/z 773.5337 and the corresponding product ion image at m/z 281.2479 and 152.9953.

e) Mobility-constrained MALDI-iPRM ion images of PG 16:0_22:6 (top), showing the precursor ion image at m/z 793.5015 and the corresponding product ion images at m/z 255.2320 and 327.3623, and of BMP 18:2_20:4 (bottom), showing the precursor ion image at m/z 793.5015 and the corresponding product ion image at m/z 279.2313 and 303.2314.

Scale bar, 2 mm. Color scale bar is shown as percentage of maximum intensity. Pixel size, 20 μm; raster size, 40 μm. Data were normalized by RMS.
